# Dual recognition of structurally unrelated mildew effectors underlies the broad-spectrum resistance of Pm3e in wheat

**DOI:** 10.1101/2025.10.26.683672

**Authors:** Lukas Kunz, Zoe Bernasconi, Matthias Heuberger, Jonatan Isaksson, Alexandros G. Sotiropoulos, Ursin Stirnemann, Jigisha Jigisha, Fabrizio Menardo, Thomas Wicker, Marion C. Müller, Beat Keller

## Abstract

Broad-spectrum resistance genes are highly valuable for sustainable crop protection, yet the molecular basis of their activity is often unknown. The *Pm3* allelic series in wheat encodes NLR receptors that recognize avirulence (AVR) effectors of wheat powdery mildew. Here, we show that near-identical *Pm3* alleles vary greatly in resistance efficacy and broadness against a global mildew isolate collection and subsequently use this model system to study the mechanisms underlying broad-spectrum resistance. We demonstrate that two alleles, *Pm3d* and *Pm3e*, provide resistance against the majority of isolates worldwide, by each recognizing two AVR effectors from powdery mildew, thereby lowering the risk of resistance breakdown. While Pm3d recognizes two closely related RNase-like AVR effectors, Pm3e detects two structurally diverse AVRs, including an effector belonging to a large, uncharacterized protein family with a novel structural fold. Using chimeric Pm3 NLRs, we identify specificity-defining polymorphisms of Pm3d and Pm3e against their diverse effector targets. Lastly, we demonstrate that Pm3d and Pm3e activities can be combined in engineered Pm3 NLRs, thereby further extending their recognition spectrum. Our findings highlight the potential of Pm3 immune receptors for long-lasting wheat protection by demonstrating their versatility in recognizing structurally diverse effectors and their amenability to NLR engineering.

## Introduction

Crop protection through breeding of genetically resistant cultivars represents a cornerstone of sustainable agriculture^1^. The underlying resistance genes frequently encode nucleotide-binding leucine rich repeat (NLR) receptors recognizing specific pathogen effectors (avirulence effectors, AVRs)^2^. AVR recognition by NLRs typically triggers a localized cell death response (hypersensitive response, HR) thereby restricting pathogen proliferation^3,4^.

Wheat is among the most widely cultivated crops worldwide^5^. However, its production is threatened by various pathogens, including wheat powdery mildew (*Blumeria graminis* f. sp. *tritici*, hereafter *Bgt*), which is considered one of the 10 biggest challenges among fungal crop pathogens^6^. Wheat breeding efforts have defined several dozen resistance genes against powdery mildew, among them several that encode NLR receptors capable of recognizing parts of the impressive effector diversity of *Bgt*^7,8^. To date, the AVRs recognized by wheat NLRs Pm1, Pm2, Pm3, Pm8, Pm17 and Pm60 have been identified^9–16^. Interestingly, all known AVR effectors in *Bgt* and the closely related barley powdery mildew (*Blumeria hordei*, *Bh*), exhibit an RNase-like structure and belong to the RNase-Like Proteins associated with Haustoria (RALPH) effector superfamily^17,18^. RALPH effectors comprise more than half of the total effector repertoire of the *Bgt* and *Bh* pathogens^17^. Despite the prevalence of RALPH effector genes in the genome, *Bgt* also encodes several hundred candidate effector proteins that do not belong to this superfamily^12^. Whether some of these effectors can also be recognized by wheat immune receptors and hence represent AVRs is currently unknown.

AVR effectors continuously evolve and diversify to evade NLR recognition, a phenomenon that is particularly pronounced in fast evolving fungal pathogens. Consequently, many NLRs are overcome quickly, resulting in race-specific resistance activity that protects crops from only a subset of pathogen races^19^. Therefore, it is of great importance to identify and molecularly characterize new resistances with broad recognition spectra that can more durably withstand pathogen evolution. Strategies to identify broadly active resistance genes include the introgression of new resistances from closely related crop species or wild crop relatives. In recent years, NLR engineering aiming at expanding the recognition spectrum through introduction of naturally occurring, or new, structure-inferred polymorphisms shows great promise to broaden NLR-based resistance^20,21^. Additionally, broadly active NLRs may exist in nature as standing genetic diversity of previously described race-specific genes. In particular, extensive allelic series of NLRs hold great promise for the identification of variants with broader recognition spectra.

The *Pm3* resistance gene in wheat comprises at least 17 functional alleles (*Pm3a*-*Pm3t*) each conferring race-specific resistance against *Bgt*. These alleles encode highly similar NLR receptors with >97% sequence identity, making *Pm3* one of the most extensive allelic series in crops^22,23^. Several allele pairs, such as *Pm3a*/*Pm3f* and *Pm3b*/*Pm3c,* were found to recognize overlapping *Bgt* race spectra by recognizing the same AVR effectors yet differ in recognition sensitivity and HR amplitude^9,11,24^. Up to now, three *AVRs* of the *Pm3* allelic series are known, the RALPH effectors *AvrPm3a/f*, *AvrPm3b/c* and *AvrPm3d*, recognized by the allelic pairs *Pm3a/Pm3f*, *Pm3b/Pm3c* and the *Pm3d* allele, respectively^9,11^. Investigations into the evolutionary origins of *Pm3* alleles have identified *Pm3CS* as the ancestral allele, which has however been completely overcome by modern *Bgt* isolates^25^. Some functional alleles (*Pm3a/Pm3b/Pm3c/Pm3f*) arose from gene conversion events with other NLR genes, resulting in highly polymorphic sequence blocks, whereas others (*Pm3d/Pm3e/Pm3g*) are nearly identical with the ancestral *Pm3CS* allele, differing by only two to five amino acids (Fig. 1a)^25,26^. The detailed study of these *Pm3CS*-like alleles has so far been hampered by the lack of near-isogenic lines that would allow to define recognition spectra without confounding genetic background effects. Moreover, previous studies tested *Pm3* alleles against local *Bgt* populations, but not against a global *Bgt* diversity panel^27^, which would be key to identify alleles with broad-spectrum resistance.

**Figure 1.**
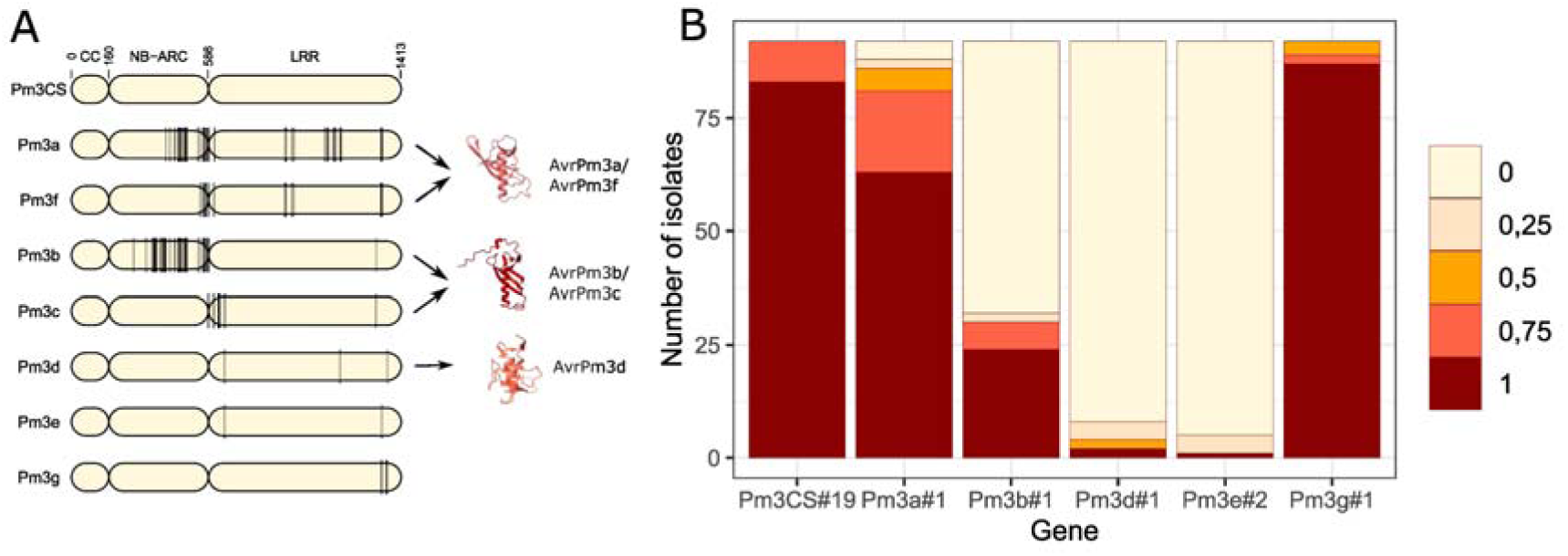
*Pm3e* and *Pm3d* provide broad-spectrum resistance against a global *Bgt* population. **A)** Schematic representation of NLR variants encoded by allelic *Pm3* genes. Black lines indicate amino acid polymorphisms compared to the ancestral, non-functional Pm3CS variant serving as a reference. AlphaFold models of the known AvrPm3a/f, AvrPm3b/c, and AvrPm3d avirulence proteins are included. Black arrows indicate recognition of the AvrPm3 proteins by their corresponding Pm3 NLR variants. **B)** Summary of the resistance response of transgenic lines expressing individual *Pm3* alleles in the wheat cultivar ‘Bobwhite’ against a global collection of 92 *Bgt* isolates. *Bgt* leaf coverage was assessed 8-9 days after infection using a scale with five categories ranging from 0 (0% leaf coverage) to 1 (100% leaf coverage).

Here, we use transgenic wheat lines to systematically compare recognition spectra of various *Pm3* alleles against 92 *Bgt* isolates collected worldwide. We show that *Pm3d* and *Pm3e* provide broad-spectrum resistance and identify their corresponding AVR effectors. We reveal that Pm3e recognizes two structurally divergent AVRs, whereas Pm3d resistance is based on the recognition of two RALPH effectors. Finally, we demonstrate that Pm3d and Pm3e resistance can be combined through NLR engineering, resulting in immune receptors that exhibit further extended recognition spectra.

## Results

### *Pm3d* and *Pm3e* confer broad-spectrum resistance against *Bgt*

To compare the resistance spectra of the *Pm3a-g* alleles in the absence of potential genetic background effects in *Pm3* donor lines, we used previously generated transgenic wheat lines expressing individual *Pm3* alleles in the susceptible wheat cultivar ‘Bobwhite’^28–31^. We investigated the resistance provided by *Pm3a*, *Pm3b*, *Pm3d*, *Pm3e*, *Pm3g* and the ancestral allele *Pm3CS,* while omitting *Pm3c* and *Pm3f* since their resistance spectra are fully covered by *Pm3b* and *Pm3a*, respectively^24^ (Fig. 1a). For each allele, we chose a single transgenic event that was previously found to exhibit high transgene expression with no pleiotropic effects^28–31^. We infected these transgenic lines with 92 *Bgt* isolates selected from a previously described global *Bgt* panel^15,27^.

Individual *Pm3* alleles exhibited markedly different resistance spectra against the global *Bgt* population ranging from no resistance to broad-spectrum resistance against nearly all tested isolates (Fig. 1b). Our data confirms that the ancestral allele *Pm3CS* has been completely overcome by *Bgt*. Similarly, *Pm3g,* which differs from *Pm3CS* by five amino acids, did not confer significant resistance (*Bgt* leaf coverage ≤0.5) against most isolates. The two highly divergent alleles *Pm3a* and *Pm3b* provided resistance against 12% and 67% of the global *Bgt* population respectively. Importantly, *Pm3d* and *Pm3e* provided broad-spectrum resistance against nearly all tested *Bgt* isolates worldwide, with only two and one virulent isolate, respectively (Fig. 1b). This finding is remarkable, given that Pm3d and Pm3e differ by only three and two amino acids from the non-functional Pm3CS (Fig. 1a).

We conclude that the group of *Pm3CS*-like alleles, namely *Pm3CS*, *Pm3d*, and *Pm3e,* in combination with their corresponding AVR factors, provide a highly interesting model system to study the molecular basis of broadly active resistance alleles. However, while a previous study has identified the RALPH effector gene *BgtE-20069b* as *AvrPm3d*^11^, the identity of the *AvrPm3e* gene is unknown.

### Avirulence on the *Pm3e* resistance allele is controlled by two *AVR* loci in *Bgt*

To identify *AvrPm3e*, we applied the *Bgt* avirulence depletion assay (AD assay), which relies on the generation of haploid F1 populations between parental isolates of opposite phenotype, followed by *R* gene-mediated selection and bulk sequencing to identify genomic regions exhibiting avirulence allele depletion (described in^16^).

We performed three crosses between isolates exhibiting opposite phenotypes on the Pm3e#2 transgenic line. Namely, we generated crosses between the *Pm3e*-avirulent isolates CHN_52-27 or IRN_GOR_2 and the isolate CHN_17-40, the only *Pm3e*-virulent isolate in our survey (Fig. 2). Additionally, we crossed CHN_52-27 to CHVD_042201, a Swiss isolate that was recently discovered to break *Pm3e* resistance in an independent study (Fig. 2)^32^. The three F1 progeny populations were then grown on the Pm3e#2 transgenic line, resulting in a *Pm3e*-selected (*i.e. AvrPm3e*-depleted) bulk, or on the susceptible wheat cultivar ‘Kanzler’, resulting in a non-selected control bulk.

**Figure 2.**
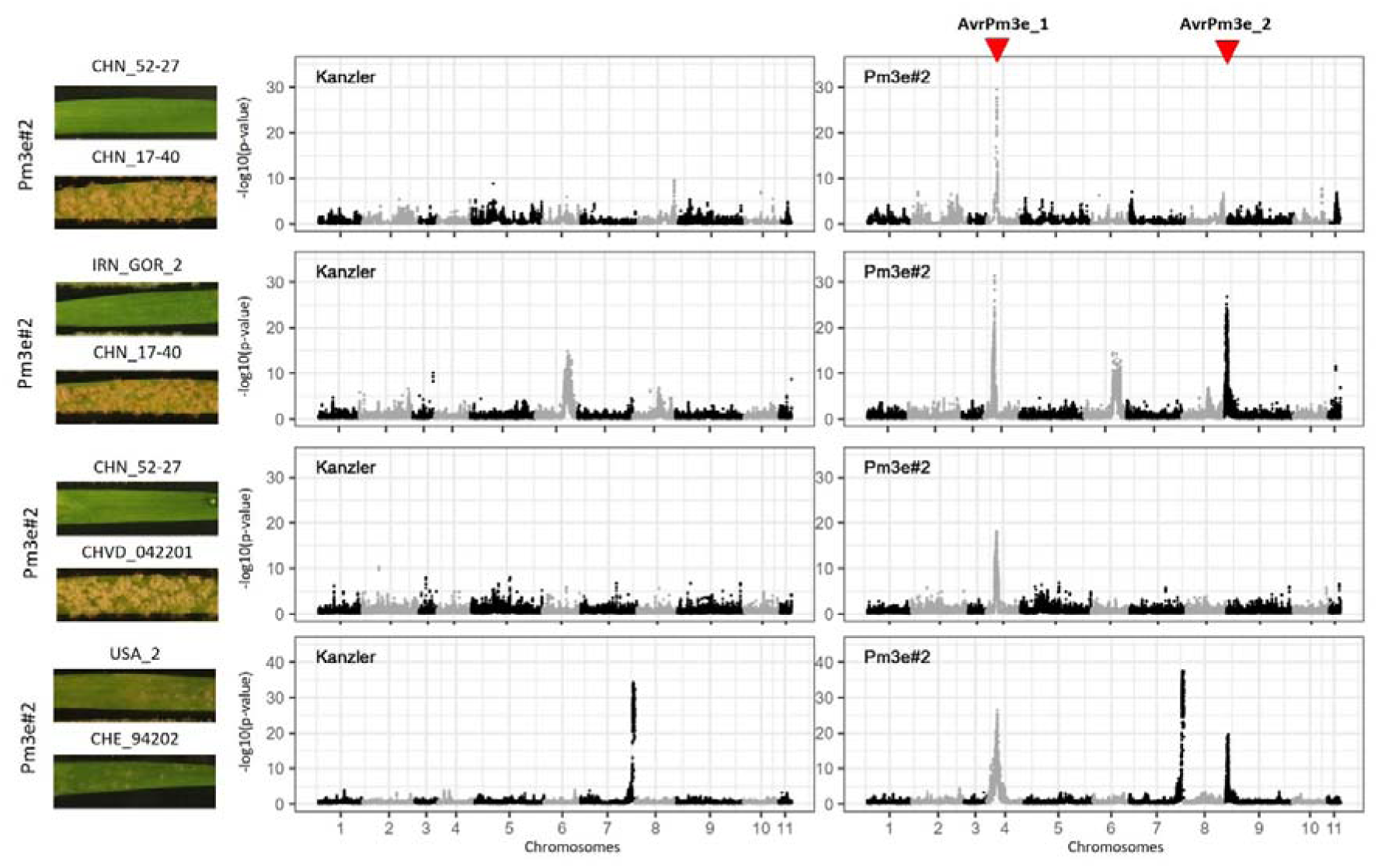
Avirulence depletion mapping in *Bgt* reveals two genetic loci associated with avirulence on *Pm3e*. Avirulence depletion mapping of *AvrPm3e* loci in four independent crosses between *Bgt* isolates. Virulence phenotypes of the parental isolates on the Pm3e#2 transgenic wheat line are shown on the left. Statistical analysis of sequenced F1 bulks grown on ‘Kanzler’ (non-selected control) or selected on the Pm3e#2 transgenic line (*Pm3e*-selected) reveal regions with deviations from an expected 1:1 parental genotype ratio, indicating avirulence allele depletion. Datapoints represent individual SNP markers along the 11 chromosomes of *Bgt* isolate CHE_96224 (x-axis) with their -log10 transformed p-values (y-axis) when testing for deviation from expected 1:1 segregation (G-test). The two genomic regions exhibiting *Pm3e*-specific depletion signals are highlighted and labelled as *AvrPm3e_1* (common to all four crosses) and *AvrPm3e_2* (only found in the IRN_GOR_2 x CHN_17-40 and the USA_2 x CHE_94202 crosses).

Additionally, we generated *Pm3e*-selected and non-selected bulks from a fourth cross between the isolates USA_2 and CHE_94202, which are both avirulent on the Pm3e#2 line (Fig. 2). This cross was initially designed for an independent project; however, we recovered viable progeny on the Pm3e#2 line, suggesting that the parental isolates USA_2 and CHE_94202 carry genetically unlinked *AvrPm3e* components.

After sequencing *Pm3e*-selected and unselected bulks of each cross, we identified regions with strong deviations from an expected 1:1 segregation ratio of parental genotypes to define regions under *Pm3e*-mediated selection, likely harboring *AvrPm3e* components (see Methods for details).

For the crosses IRN_GOR_2 x CHN_17-40 and USA_2 x CHE_94202, our non-selected control bulks revealed one region in each cross, that deviated from the expected 1:1 ratio of parental genotypes even in the absence of *Pm3e*-mediated selection (Fig. 2). Such effects are likely caused by the presence of specific virulence factors or due to genetic incompatibility between parental isolates in the corresponding genomic regions (for details see Sup Note S1). To ensure the identification of *Pm3e*-specific selection signals, we excluded these regions from further analysis.

Focusing on *Pm3e*-specific signals of selection, we identified a candidate locus on Chr-04 for all four crosses, which we defined as the *AvrPm3e_1* locus. Additionally, we identified a second *Pm3e*-specific peak on Chr-09 in two of the four crosses which we designated as the *AvrPm3e_2* locus (Fig. 2). In the USA_2 × CHE_94202 cross, the *Pm3e*-selected bulk showed depletion of CHE_94202 at the *AvrPm3e_1* locus and depletion of USA_2 at the *AvrPm3e_2* locus, indicating that each parental isolate carries a genetically distinct *AvrPm3e* component. Taken together, our data revealed that avirulence towards *Pm3e* in *Bgt* is likely controlled by two independently segregating *AVR* loci.

### AvrPm3e_1 is a novel type of avirulence effector

To functionally validate AvrPm3e candidate effectors, we first focused on the *AvrPm3e_1* locus on Chr-04, common to all four crosses. We defined the genomic interval exhibiting the strongest selection signal for each cross (see methods) and subsequently defined the overlap of these intervals based on the *Bgt* reference assembly of *Pm3e*-avirulent isolate CHE_96224 (Bgt_genome_v3_16^33^ (Sup Table 1). The overlapping interval had a size of 3.2 kb harboring a single gene, *BgtE-5754*, encoding an effector protein with 367 amino acids which is significantly larger than previously identified AVRs in the *Blumeria* genus (Fig. 3a). Importantly, the *BgtE-5754* gene exhibits non-synonymous polymorphisms between all respective parents of the four genetic crosses, making it a prime *AvrPm3e_1* candidate (Fig. 3b, Sup Fig. 3a,d,g).

**Figure 3.**
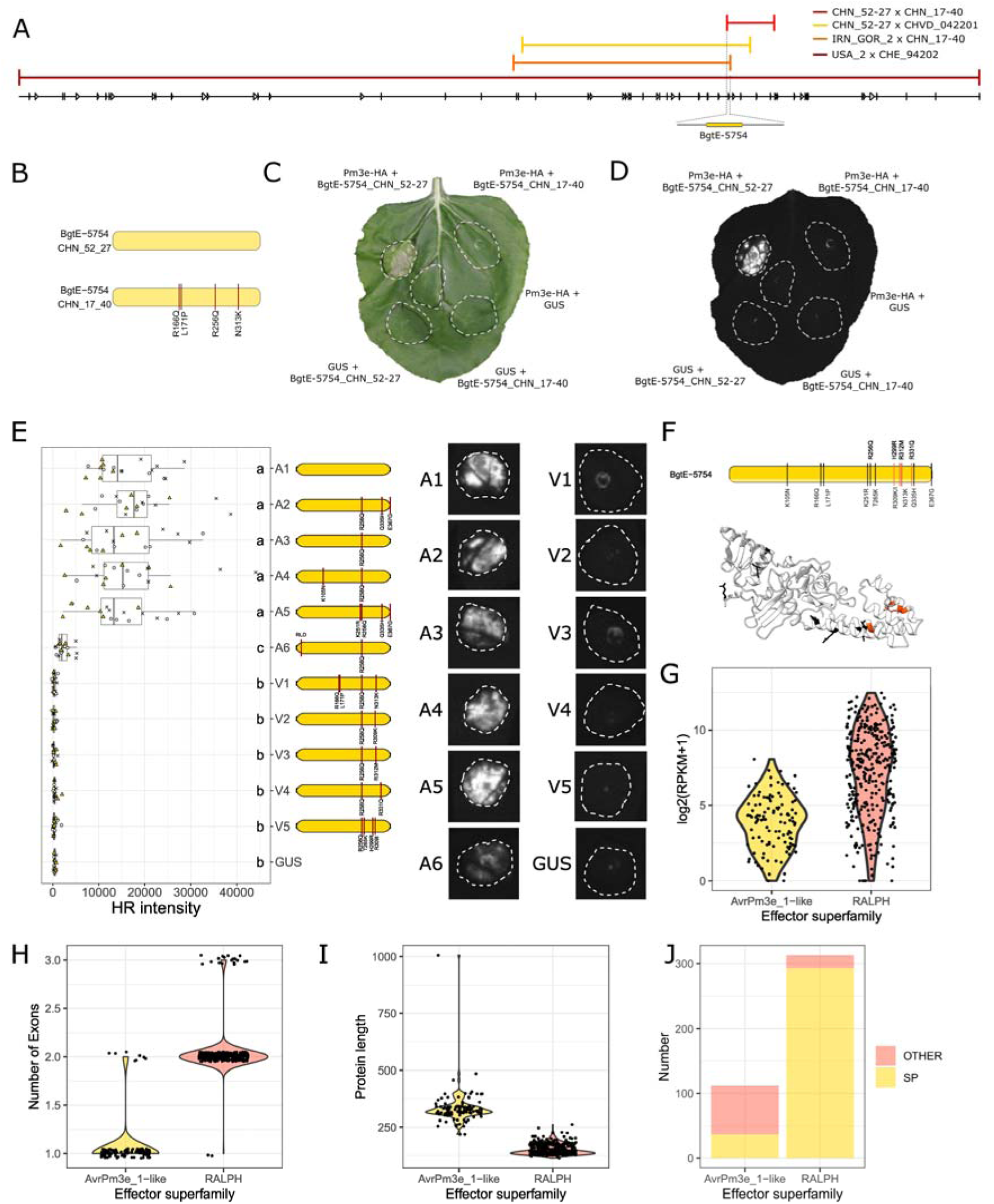
AvrPm3e_1 (BgtE-5754) belongs to a structurally novel family of *Bgt* effectors, distinct from RALPH effector proteins. **A)** A single genomic region on chromosome 04 exhibits a *Pm3e*-specific depletion signal in all four genetic crosses. Genomic regions with the strongest depletion signal (see Fig. 2) for each cross are shown and aligned to the CHE_96224 reference genome. The *AvrPm3e_1* candidate locus, defined by the overlap between all four crosses, is shown as a zoom-in and contains a single effector gene, *BgtE-5754* (yellow gene model), which exhibits non-synonymous polymorphisms between the parental isolates of all four crosses. **B)** Structural model of the AvrPm3e_1 candidate effector BgtE-5754 in the *Pm3e*-avirulent isolate CHN_52-27 and the virulent isolate CHN_17-40, with four non-synonymous polymorphisms indicated by red lines. **C & D)** *Agrobacterium*-mediated co-expression of BgtE-5754_CHN_52-27 (BgtE-5754-A1) or BgtE-5754_CHN_17-40 (BgtE-5754-V1) with Pm3e-HA or a GUS negative control in *N. benthamiana*. Leaves were imaged with a standard camera (C) or a Fusion FX imager (D). The experiment was repeated three times with n = 6 leaves per experiment (total n = 18). **E)** *Agrobacterium*-mediated co-expression of Pm3e-HA with various BgtE-5754 (AvrPm3e_1) variants or GUS. Boxplots show quantified HR intensities from 18 infiltration spots per combination, obtained from three independent experiments (denoted by distinct symbols). Letters above boxplots denote statistically significant differences at p < 0.05 based on Dunn’s test. Representative pictures for each combination are shown on the right. **F)** Schematic representation of the BgtE-5754 effector protein showing all naturally occurring amino acid polymorphisms found in *Bgt* compared to the BgtE-5754-A1 variant. AlphaFold3 structural prediction of BgtE-5754 is shown below. Model quality scores are provided in Sup Fig. 5. Polymorphisms in black were found to have no effect on *Pm3e* recognition, while polymorphisms highlighted in red disrupt recognition. **G–J)** Comparison between proteins encoded by *AvrPm3e_1*-like genes/proteins and those belonging to the RALPH effector family. Datapoints represent individual effectors with data point shapes indicating their effector family according to nomenclature of Müller et al. (2019)^33^. **G)** Protein length of all family members shown as violin plots. **H)** Number of exons per gene displayed as violin plots. **I)** Effector expression levels in reference isolate CHE_96224 at two days post-inoculation on the susceptible wheat line ‘Chinese Spring’. Data points represent log_₂_(RPKM + 1) values, shown as violin plots. **J)** Number of proteins predicted to have a signal peptide in each effector superfamily (according to SignalP6.0).

Signal peptide (SP) prediction using different software was inconsistent for BgtE-5754 (see Sup Fig. 1, Sup Note S2). Therefore, we co-expressed the entire coding sequence of *BgtE-5754* with *Pm3e* in *Nicotiana benthamiana* and assessed its ability to elicit a *Pm3e*-specific hypersensitive response (HR). First, we focused on the haplovariants identified in the parental isolates CHN_52-27 (*Pm3e*-avirulent) and CHN_17-40 (virulent). Indeed, the BgtE-5754_CHN_52-27 variant (identical to CHE_96224 reference) elicited strong Pm3e-dependent HR, whereas the variant from the virulent parent CHN_17-40, differing by four amino acids (R166Q, L171P, R256Q, N313K), did not (Fig. 3b-d). Importantly, neither of the two BgtE-5754 variants, nor Pm3e, induced an HR response upon co-expression with a GUS negative control (Fig. 3c-d). We therefore concluded that *BgtE-5754* is indeed *AvrPm3e_1* and designated the avirulent BgtE-5754_CHN_52-27 variant as BgtE-5754-A1 and the virulent BgtE-5754_CHN_17_40 variant as BgtE-5754-V1. Of note, expression of N-terminally truncated versions of BgtE-5754-A1, corresponding to different putative signal peptide predictions, did not result in Pm3e-mediated HR, suggesting that full-length BgtE-5754 acts as AvrPm3e_1 (Sup Fig. 1, Sup. Note S2).

For the remaining crosses, the genetic analysis suggested that IRN_GOR_2 and CHE_94202 would carry recognized variants of BgtE-5754. BgtE-5754_IRN_GOR_2 (hereafter BgtE-5754-A2) and BgtE-5754_CHE_94202 (BgtE-5754-A3) differ by three and one amino acids from BgtE-5754-A1, respectively (R256Q, Q335H, E367G in IRN_GOR_2; R256Q in CHE_94202). Both variants triggered strong HR upon co-expression with *Pm3e* (Sup Fig. 2). Conversely, the parental isolates CHVD_042201 and USA_2 are predicted to contain non-recognized variants of BgtE-5754. In both isolates, BgtE-5754 differs by two amino acid polymorphisms from BgtE-5754-A1 (R256Q, R309K in CHVD_042201; R256Q, R312M in USA_2), consequentially named BgtE-5754-V2 and V3. In agreement with our genetic analysis neither BgtE-5754-V2 nor BgtE-5754-V3 elicited a Pm3e-mediated HR response (Sup Fig. 2). We therefore concluded that *BgtE-5754* represents the *AvrPm3e* component responsible for the mapped *AvrPm3e_1* region in all four *crosses*.

To ensure that the lack of recognition of virulent BgtE-5754 variants was not due to an absence of protein production in *N. benthamiana*, we tagged representative avirulent and virulent variants BgtE-5754-A1, A2, V1 and V2 with C-terminal mRFP-FLAG epitopes. Western blot analysis revealed that all four tested variants were efficiently produced in *N. benthamiana*, indicating that amino acid polymorphisms in the virulent variants BgtE-5754-V1 and V2 do not negatively affect protein amounts, but are rather involved in Pm3e evasion (Sup Fig 3). Interestingly, the western blot analysis also revealed that BgtE-5754 might undergo posttranslational processing in *N. benthamiana*, although it is currently unclear whether this phenomenon is biologically relevant (see Sup Note S3).

In addition to the six BgtE-5754 variants identified in the parental isolate used for mapping, we detected five additional variants among 220 global *Bgt* isolates^27^. Of these, three triggered a Pm3e-dependent HR (BgtE-5754-A4, A5, A6), whereas the remaining two did not (BgtE-5754-V4, V5) (Fig. 3e, Sup Note S4). To identify the polymorphisms responsible for Pm3e evasion, we tested consequences of individual amino acid substitutions in the BgtE-5754-A1 backbone, revealing that all naturally occurring polymorphisms that evade Pm3e-mediated recognition are located in a short C-terminal stretch of the BgtE-5754 protein (Fig. 3f, Sup Note S4, Sup Fig. 4). In the absence of structural information for BgtE-5754 or any closely related effectors, we used AlphaFold to predict its structure (Sup Fig 5). This revealed that BgtE-5754 is structurally unrelated to the RNase-like fold found in all previously identified AVR effectors in *Bgt*, belonging to the RALPH effector superfamily (Fig. 3f). Pfam analysis did not reveal any strong homology to known proteins. Interestingly, four of the immune-evasive amino acid polymorphisms affect three positions within the same predicted alpha helix in the BgtE-5754 structure, while the fifth affects the neighboring alpha helix (Fig. 3f), indicating this C-terminal domain is involved in BgtE-5754 recognition.

**Figure 4.**
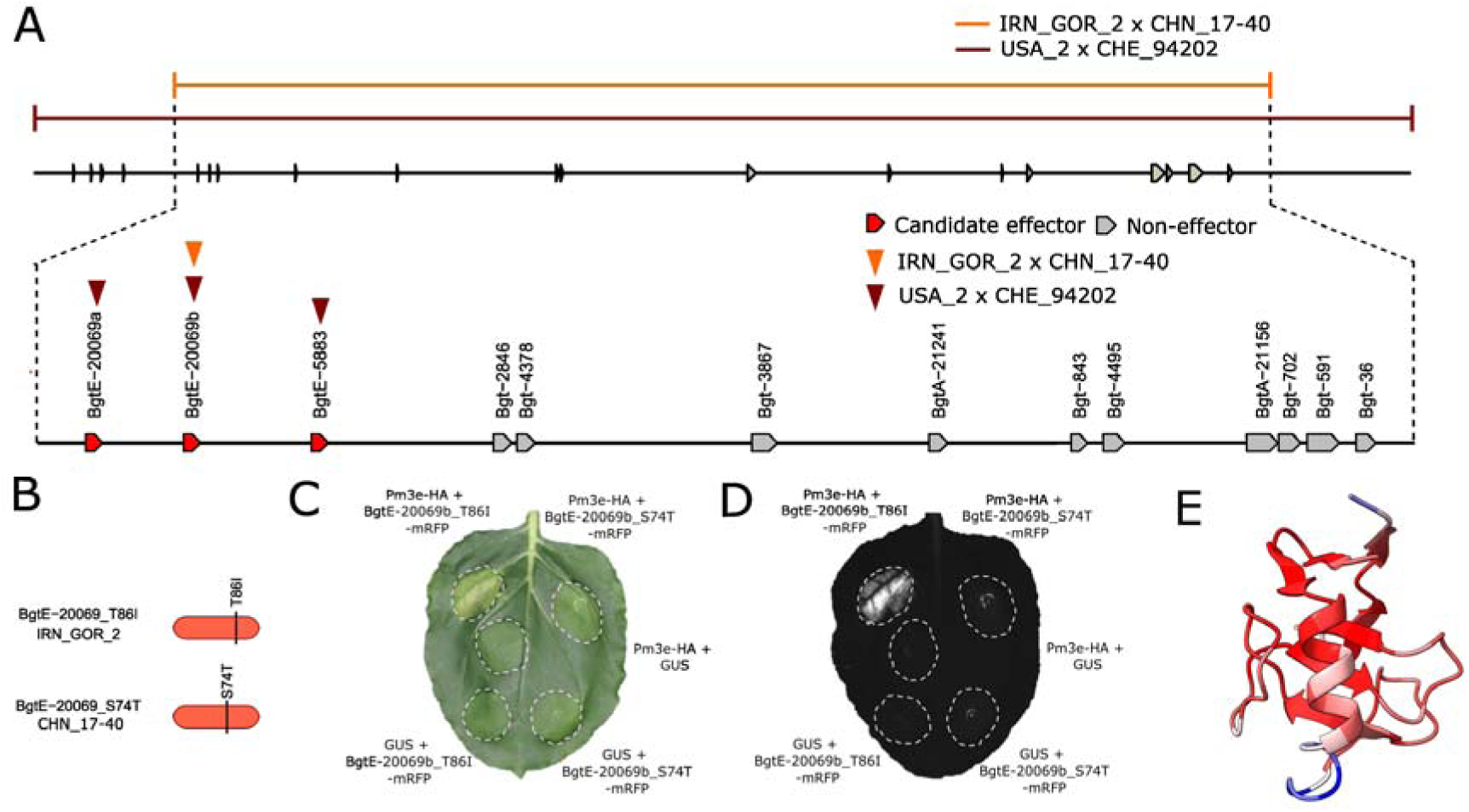
AvrPm3e_2 (BgtE-20069b) is a RALPH effector. **(A)** Genomic region on chromosome 09 exhibiting a *Pm3e*-specific depletion signal in two out of four crosses (IRN_GOR_2 x CHN_17-40 and USA_2 x CHE_94202). Regions with the strongest depletion signal for both crosses are shown and aligned to the CHE_96224 reference genome. The *AvrPm3e_2* candidate locus, defined by the overlap between both crosses, in the reference genome CHE_96224 is shown as a zoom-in. Effector genes (red) and non-effector genes (grey) are represented by arrows. Gene models are not drawn to scale. Effectors exhibiting non-synonymous polymorphisms in the parental isolates of individual crosses are marked by orange (IRN_GOR-2 x CHN_17-40) and dark red (USA_2 x CHE_94202) arrowheads. **(B)** Schematic protein model of the AvrPm3e_2 candidate effector BgtE-20069b in the *Pm3e*-avirulent isolate IRN_GOR-2 and virulent isolate CHN_17-40. Sequence polymorphisms are defined in comparison with BgtE-20069b sequence in the CHE_96224 reference isolate and are highlighted by black lines. **(C & D)** *Agrobacterium*-mediated co-expression of BgtE-20069b_T86I-mRFP or BgtE-20069b_S74T-mRFP with Pm3e-HA or a GUS negative control in *N. benthamiana*. Leaves were imaged with a camera (C) or a Fusion FX imager (D). The experiment was performed three times with n=6 leaves per experiment (total n=18). **(E)** Predicted 3D structure of BgtE-20069b_T86I according to Alphafold3 modelling. Coloring represents predicted local distance difference test (pLDDT) local confidence according to Alphafold (100 (dark red) – 0 (dark blue)).

**Figure 5.**
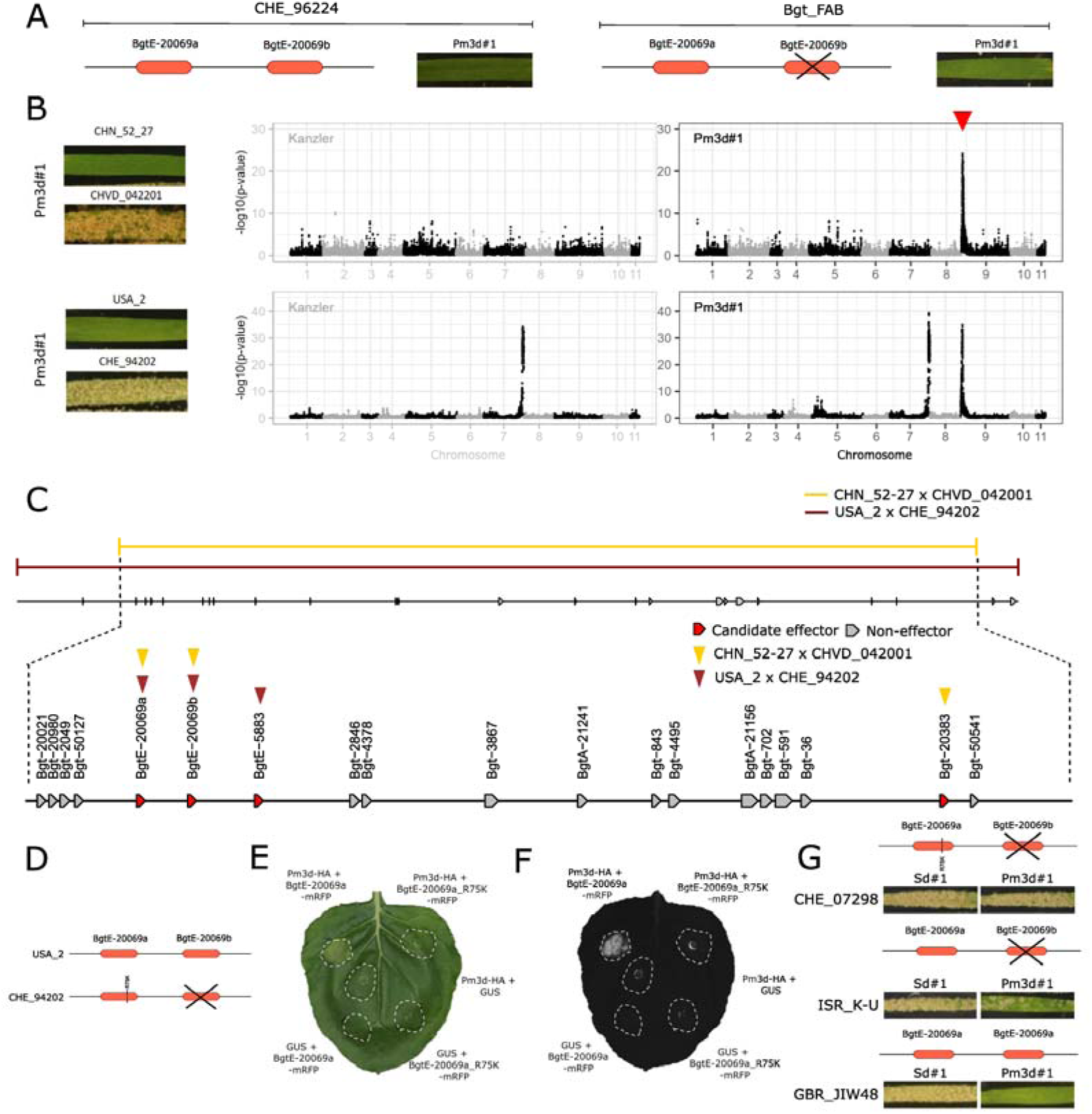
Avirulence on *Pm3d* is controlled by two neighboring *AVR* genes encoding RALPH effector proteins. **(A)** The UV mutant Bgt_FAB exhibits an avirulent phenotype on the Pm3d#1 transgenic wheat line, despite carrying a deletion of the *BgtE-20069b* (*AvrPm3d_1/AvrPm3e_2*) effector gene. The phenotype of wild type *Bgt* isolate CHE_96224, used for UV mutagenesis, is shown as comparison. **(B)** Avirulence depletion mapping on *Pm3d* in two independent *Bgt* crosses. Virulence phenotypes of the parental isolates on the Pm3d#1 transgenic wheat line are shown on the left. Statistical analysis of sequenced F1 bulks selected on ‘Kanzler’ (non-selected control) or the Pm3d#1 transgenic line (Pm3d-selected) are shown on the right. ‘Kanzler’ bulks are drawn with grey lines to represent that these panels were reprinted from Fig. 2. Datapoints represent individual SNP markers along the 11 chromosomes of *Bgt* isolate CHE_96224 (x-axis) with their -log10 transformed p-values (y-axis) when testing for deviation from expected 1:1 segregation (G-test). *Pm3d*-specific depletion signal is indicated by a red arrowhead. **(C)** The *AvrPm3d* candidate locus, defined by the overlap between both crosses, in the reference genome CHE_96224. Effector genes (red) and non-effector genes (grey) are represented by arrows. Gene models are not drawn to scale. Effectors exhibiting non-synonymous polymorphisms in the parental isolates of individual crosses are marked by yellow (CHN_52-27 x CHVD_042001) and dark red (USA_2 x CHE_94202) arrowheads. **(D)** Gene models of the neighboring effector orthologs *BgtE-20069a* and *BgtE-20069b* in the *Pm3d*-avirulent isolate USA_2 and virulent isolate CHE_94202. The non-synonymous mutation (leading to a R75K substitution in *BgtE-20069a)* is highlighted by a red vertical line, the deletion of *BgtE-20069b* in CHE_94202 is indicated by crossed black lines. **(E & F)** *Agrobacterium*-mediated co-expression of BgtE-20069a-mRFP (AvrPm3d_2) or BgtE-20069a_R75K-mRFP with Pm3d-HA or a GUS negative control in *N. benthamiana*. Leaves were imaged with a camera (E) or a Fusion FX imager (F). The experiment was performed three times with n=6 leaves per experiment (total n=18). **(G)** Representative phenotyping pictures of isolates that carry a *BgtE-20069b* deletion on the Pm3#d1 transgenic line and the corresponding sister line Sd#1. For each isolate, a schematic representation of the two genes is depicted. A black cross indicates *BgtE-20069b* deletion.

The observation that BgtE-5754 is significantly larger than previously identified AvrPm3 effectors and appears structurally unrelated to RALPH effectors prompted us to investigate this effector structure in more detail. Based on Alphafold2 structural modelling and TM-align to infer structural similarity, we found 112 proteins exhibiting structural similarity with BgtE-5754 (AvrPm3e_1) in *Bgt*, which we consequentially refer to as the AvrPm3e_1-like superfamily (for details see Sup Note S5). This indicates that AvrPm3e_1-like effectors, comparable to the RALPH superfamily, constitute an expanded and diversified effector superfamily that constitutes ∼13% of the effectors in *Bgt* (Sup Note S5). RALPH effector genes have been previously described to exhibit high average expression levels at early infection stages and to consist of two protein-coding exons resulting in small effector proteins (100-150 residues) that predominantly contain N-terminal signal peptides for export via the secretory pathway (Fig. 3g-j,^17,18^). In contrast, we found *AvrPm3e_1*-like effector genes to exhibit lower average expression during haustorial stages and to predominantly consist of a single large exon that gives rise to much larger effector proteins (∼300-400 residues) (Fig. 3g-i). Additionally, we found that AvrPm3e_1-like effectors would often lack a classic signal peptide (Fig. 3j), as exemplified by AvrPm3e_1 (BgtE-5754) itself, indicating they might be secreted by non-conventional pathways (Sup Fig. 1, Sup Note S2).

Based on structural predictions from a previous study^34^ and publicly available databases, we investigated the effector repertoire of *Bgt* sister species *Bh* and other more divergent powdery mildew species. This analysis identified 82 AvrPm3e_1-like effector proteins in *Bh*, whereas none were identified in other powdery mildew species, suggesting that this effector group is specific to the *Blumeria* genus (see Sup Note S5).

### *Pm3e* recognizes a second AVR effector, AvrPm3e_2, with an RNase-like structure

In a next step, we investigated the *AvrPm3e_2* locus on Chr-09 for the presence of an additional *AVR* factor (Fig. 2). This locus was identified in two out of the four performed genetic crosses (IRN_GOR_2 x CHN_17-40 and USA2 x CHE_94202). We defined the *AvrPm3e_2* locus based on the overlapping genomic regions with strongest selection signals for both crosses based on the reference assembly of CHE_96224 (Bgt_genome_v3_16) (Fig 4a). The identified interval of 312 kb contains 13 high-quality annotated genes including three effector candidates. Importantly, a single effector candidate (*BgtE-20069b* in CHE_96224) is polymorphic between parental isolates of both crosses and was therefore considered the best *AvrPm3e_2* candidate (Fig 4a, b). Between IRN_GOR_2 and CHN_17-40, the BgtE-20069b effector differs by two amino acid polymorphisms, BgtE-20069b_T86I in IRN_GOR_2 (using *BgtE-20069b* in CHE_96224 as a reference) and BgtE-20069b_S74T in CHN_17-40 (Fig 4b). In the second cross, the *BgtE-20069b* sequence found in USA_2 is identical with the reference assembly of CHE_96224, whereas the effector gene is deleted in CHE_94202 (Sup Fig 6a).

**Figure 6.**
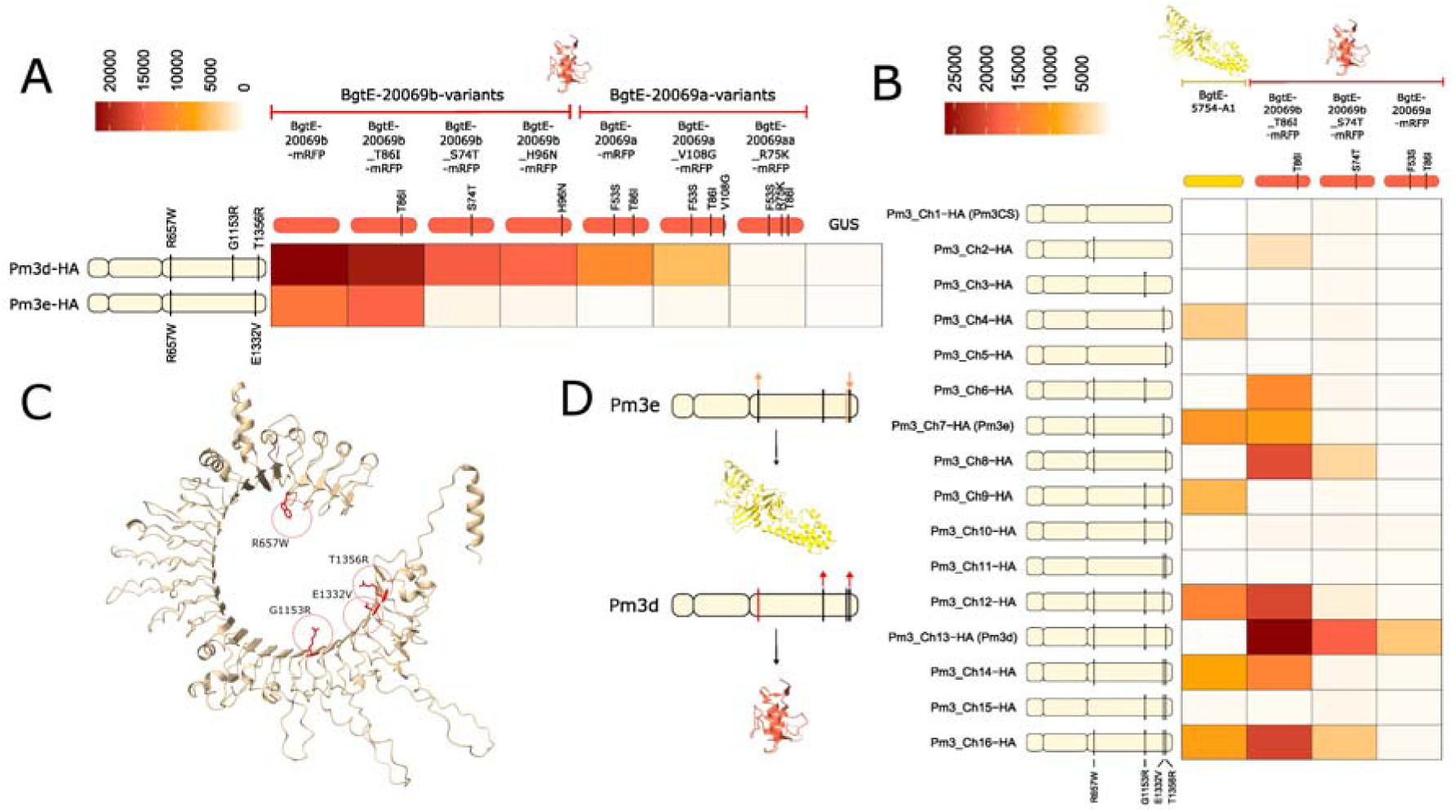
Single amino acid polymorphisms in the LRR domain of Pm3d and Pm3e determine recognition specificities towards their AVR effectors. **A)** Recognition strength of Pm3d-HA and Pm3e-HA towards variants of BgtE-20069b-mRFP (AvrPm3d_1/AvrPm3e_2) and BgtE-20069a-mRFP (AvrPm3d_2) in *N. benthamiana*, shown as a heat map. Schematic representations of amino acid polymorphisms defining Pm3d and Pm3e are shown on the left. Schematic representations of BgtE-20069b/BgtE-20069a variants with defining amino acid polymorphisms relative to the BgtE-20069b variant in reference isolate CHE_96224 (left) are shown above the heat map. The heatmap shows the mean HR strength from n=18 individual replicates across three independent experiments (full dataset is shown in Sup Fig. 11). **B)** Heatmap depicting recognition strength of 16 C-terminally HA-tagged Pm3CS/Pm3d/Pm3e NLR chimeras towards BgtE-5754-A1 (AvrPm3e_1) and selected C-terminally mRFP-tagged BgtE-20069b/BgtE-20069a variants. Schematic representations of amino acid polymorphisms defining the individual chimeras are shown on the right. Amino acid polymorphism combinations are illustrated in the drawing relative to the Pm3CS reference. The heatmap shows the mean HR strength from n=18 individual replicates across three independent experiments (full dataset is shown in Sup Fig. 14). **C)** AlphaFold3 structural prediction of the LRR domain of the Pm3_Ch16 construct combining all Pm3d-/Pm3e-specific polymorphisms (highlighted in orange). **D)** Schematic summary of datasets depicted in (B) and Sup Fig. 14, highlighting Pm3d/Pm3e-specific polymorphisms that positively or negatively influence recognition of BgtE-5754 (top) or BgtE-20069b/BgtE-20069a (bottom). Polymorphisms highlighted by a yellow or red line are required for recognition of BgtE-5754 and BgtE-20069b/BgtE-20069a, respectively. Polymorphisms that quantitatively influence recognition strength are indicated by arrows (upward arrow = increase HR; downward arrow = decrease HR).

A previous study has identified BgtE-20069b as AvrPm3d and showed that it triggers a strong Pm3d-dependent HR in *N. benthamiana*^11^. The study also investigated possible recognition of BgtE-20069b by other Pm3 NLRs and concluded that BgtE-20069b is not recognized by Pm3e^11^. Based on our genetic indications, we tested BgtE-20069b_T86I present in the avirulent parental isolate IRN_GOR_2 and BgtE-20069b_S74T from the *Pm3e*-virulent isolate CHN_17-40 for Pm3e-mediated HR in *N. benthamiana*. Consistent with the previous study^11^, we did not observe an HR response with any of the tested BgtE-20069b variants in combination with Pm3e, whereas BgtE-20069b_T86I was readily recognized by Pm3d (Sup Fig 7). In the same study, it was observed that the BgtE-20069b protein in *N. benthamiana* was undetectable on Western blots, indicating that the effector protein might be unstable *in planta*, despite triggering HR upon co-expression with Pm3d^11^. Based on this observation, we attempted to stabilize BgtE-20069b by a C-terminal fusion of the effector with a highly soluble mRFP fluorophore. Indeed, co-expression of stabilized BgtE-20069b_T86I-mRFP with Pm3e resulted in an HR response, whereas BgtE-20069b_S74T-mRFP found in the virulent parental isolate CHN_17-40 did not (Fig 4c-d). Similarly, BgtE-20069b-mRFP from the avirulent isolate USA_2 of the second cross induced a Pm3e-mediated HR response (Sup Fig 6b,c). Therefore, we concluded that BgtE-20069b, in addition to representing AvrPm3d, is indeed AvrPm3e_2 and that the lack of previous recognition of this effector by Pm3e in the *N. benthamiana* experimental system can likely be attributed to a lack of protein stability. Thus, BgtE-20069b is an AVR for both Pm3d and Pm3e, but the two NLRs exhibit differences in sensitivity towards this effector. Owing to this complexity, we will refer to this AVR as BgtE-20069b instead of AvrPm3d/AvrPm3e_2 and specify the exact protein variants of BgtE-20069b throughout the manuscript.

**Figure 7.**
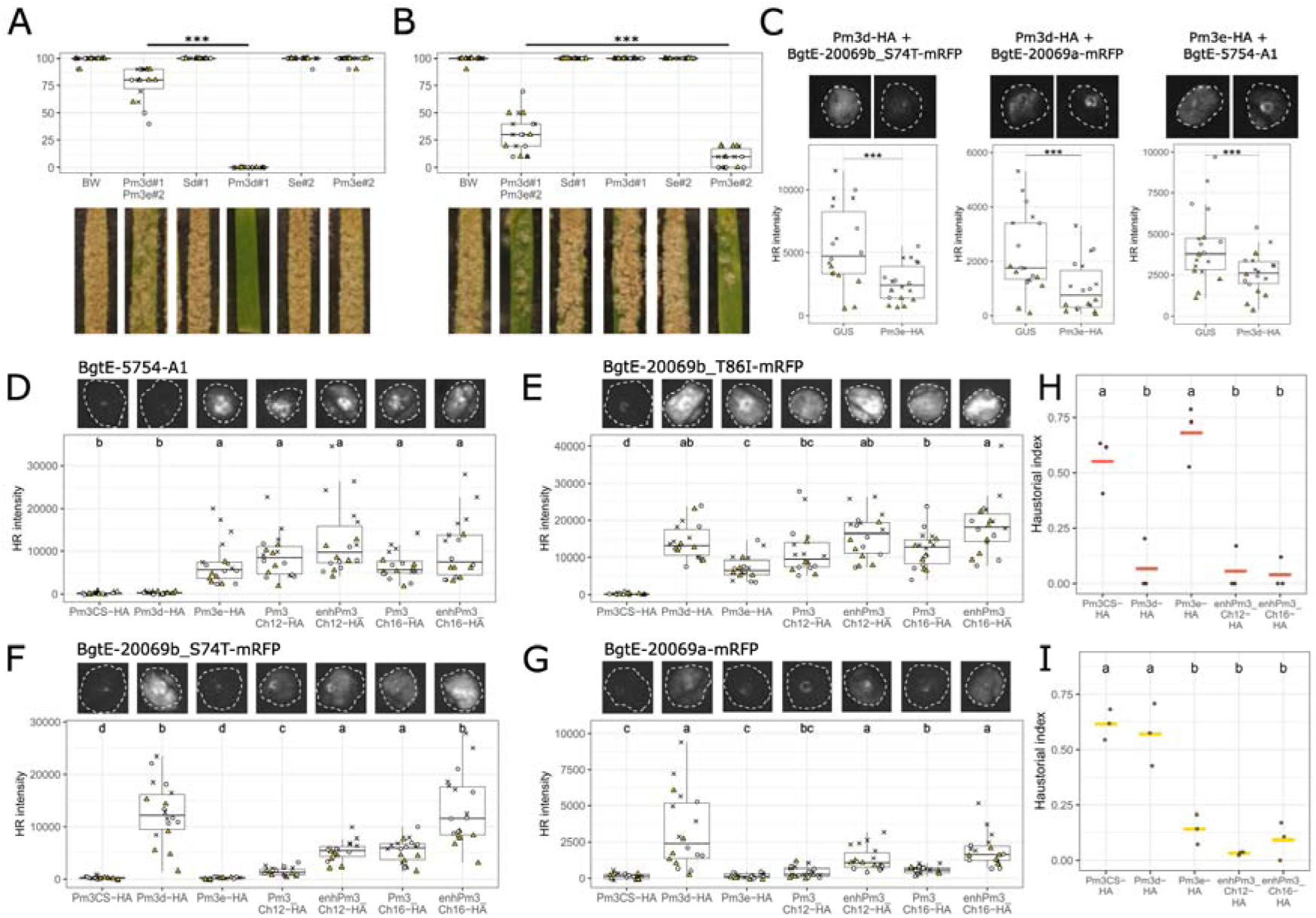
Pm3d and Pm3e recognition specificity can be combined in chimeric Pm3 NLRs. **A & B)** Virulence phenotypes of isolates CHN_17_40 (A) and CHE_94202 (B) on various wheat lines. “BW” refers to the wheat cultivar ‘Bobwhite’ used as a background for Pm3d#1 and Pm3e#2 transgenic lines. “Pm3d#1 / Pm3e#2” denotes F1 plants obtained from a cross between transgenic lines Pm3d#1 and Pm3e#2. Sd#1 and Se#2 refer to transgene-free sister lines of Pm3d#1 and Pm3e#2. Boxplots display infection scores based on a 10-step scale ranging from 0% to 100% leaf coverage, assessed 11 days post-infection (11 dpi). A total of 18 individual leaf segments were scored across three independent infection plates, indicated by different symbols. Statistical significance between the Pm3d#1 x Pm3e#2 F1 individuals and the respective parental lines was assessed using the Wilcoxon rank-sum test; *** indicates p < 0.001. **C)** Results of allelic suppression testing in *N. benthamiana* using the effector variants BgtE-20069b_S4T-mRFP, BgtE-20069a-mRFP (both exclusively recognized by Pm3d-HA) and BgtE-5754-A1 (exclusively recognized by Pm3e-HA). Boxplots show quantified HR intensities from 18 infiltration spots per combination, obtained from three independent experiments (denoted by distinct symbols). Statistical significance was assessed using a paired Wilcoxon signed-rank test; *** indicates p < 0.001. **(D - F)** *Agrobacterium-*mediated co-expression of **D)** BgtE-5754-A1 (AvrPm3e_1), **E)** BgtE-20069b_T86I-mRFP, **F)** BgtE-20069b_S74T-mRFP, and **G)** BgtE-20069a-mRFP with various Pm3-HA variants. Boxplots show quantified HR intensities from 18 infiltration spots per combination, obtained from three independent experiments (denoted by distinct symbols). Letters above boxplots denote statistically significant differences at p < 0.05 based on Dunn’s test. Representative HR images are shown above the boxplots. (**H & I)** Haustorial index in epidermal cells of wheat cultivar ‘Bobwhite’ transformed via particle bombardment with the reporter plasmid *pUbi:GUS* and the indicated *Pm3* constructs. Haustorial index represents the percentage of spores successfully forming haustoria in GUS-positive cells, assessed 44 hours post-inoculation. **H)** Haustorial index measured for *Pm3d*-avirulent, *Pm3e*-virulent isolate CHN_17_40. **I)** Haustorial index measured for *Pm3e*-avirulent, *Pm3d*-virulent isolate CHE_94202. Black dots represent individual experiments; colored lines indicate the mean across three experiments. Different letters indicate statistically significant differences according to ANOVA followed by Tukey HSD test (p < 0.05).

Importantly, BgtE-20069b is a RALPH effector with a predicted RNase-like structure (Fig. 4e,^11,17^). We conclude that Pm3e recognizes two structurally unrelated avirulence proteins, thereby explaining the broad-spectrum resistance activity of this *Pm3* allele against the global *Bgt* diversity.

### *Pm3d* recognizes two RNase-like effectors

In our survey, *Pm3d* also exhibited broad resistance activity, comparable to *Pm3e* (Fig. 1b). This raised the question whether *Pm3d* also recognizes multiple *AVRs*. Bourras et al. (2019) showed that *BgtE-20069b* in *Bgt* reference isolate CHE_96224 is recognized by *Pm3d* and that *Pm3d*-virulent isolates, like CHE_94202, lack *BgtE-20069b* due to a gene deletion^11^. However, a UV mutant of the *Pm3d*-avirulent isolate CHE_96224, termed Bgt_FAB, which was generated in our lab as part of an independent project^35^ coincidentally contains a deletion of the entire coding sequence of *BgtE-20069b* but exhibits an avirulent phenotype on the Pm3d#1 transgenic line (Fig. 5a, Sup Table 2). This strongly supports the existence of a second *AvrPm3d* component (hereafter *AvrPm3d_2*).

Since BgtE-5754 (*AvrPm3e_1*) does not trigger *Pm3d*-mediated HR in *N. benthamiana* (Sup Fig. 8), we hypothesized that *AvrPm3d_2* is represented by a third, unknown effector. To identify *AvrPm3d_2,* we performed the avirulence depletion assay using two of the four *Bgt* crosses, namely USA_2 x CHE_94202 and CHN_52-27 x CHVD_042201 for which parental isolates exhibit opposite phenotypes on the Pm3d#1 transgenic line (Fig. 5b). We identified a single *Pm3d*-specific locus on Chr-09 for both crosses (Fig 5c), which corresponded to the *AvrPm3d_1/AvrPm3e_2* locus (Fig 2). These results confirm that *AvrPm3e_1* (*BgtE-5754*), residing on Chr-04, is a *Pm3e*-specific avirulence gene and suggest that the unknown *AvrPm3d_2* component is genetically closely linked to *AvrPm3d_1/AvrPm3e_2* (*BgtE-20069b*). We therefore defined the overlapping intervals from the two crosses in the *Bgt* reference assembly of CHE_96224. Within the resulting 453 kb interval, only two effector genes were polymorphic between the parental isolates of both crosses: *BgtE-20069b*, which is deleted in the virulent parental isolates CHVD_042201 and CHE_94202, and its directly adjacent paralog *BgtE-20069a* which exhibits amino acid polymorphisms between the parents of both crosses (Fig. 5c,d, Sup Fig. 9). In a first step, we confirmed that both BgtE-20069b variants found in both avirulent isolates USA_2 (BgtE-20069b) and CHN_52-27 (BgtE-20069b_H96N) induced a Pm3d-mediated HR therefore replicating the findings of Bourras at al. (2019)^11^ and confirming BgtE-20069b as AvrPm3d_1 in both crosses (Sup Fig. 9). Next, we investigated if BgtE-20069a is AvrPm3d_2. In the reference isolate CHE_96224 and in the avirulent parental isolate USA_2, the two paralogs BgtE-20069b and BgtE-20069a differ by two polymorphisms (F53S and T86I). Due to their similarity, BgtE-20069a has previously been tested for recognition by Pm3d in *N. benthamiana,* however, no HR response was observed^11^. Given our observation that the BgtE-20069b effector can be stabilized by fusion to an mRFP fluorophore we followed a similar strategy for BgtE-20069a. Co-expression of BgtE-20069a-mRFP from avirulent isolate USA_2 with Pm3d in *N. benthamiana* triggered HR, whereas the R75K variant from virulent parent CHE_94202 (BgtE-20069a_R75K-mRFP) did not (Fig. 5e-f). Similarly, BgtE-20069a_V108G-mRFP from avirulent isolate CHN_52-27 was recognized by Pm3d, while the virulent parent CHVD_042201 also carried the immune evasive R75K variant (Sup Fig. 9d). Thus, we concluded that BgtE-20069a is AvrPm3d_2 and that R75K represents a gain-of-virulence mutation. We verified protein production of the Pm3d-immune evasive BgtE-20069a_R75K, along the recognized variants BgtE-20069b_T86I, BgtE-20069b_S74T and BgtE-20069a, via C-terminal mRFP-FLAG epitope tagging, followed by Western blot analysis. All four variants were efficiently produced to similar levels in *N. benthamiana,* indicating that the R75K gain-of-virulence mutation does not affect effector stability, but rather interferes with the Pm3d recognition (Sup Fig. 10).

Based on mapping coverage^33^ we identified 12 among 220 analyzed global isolates, that carry a *BgtE-20069b* deletion (Sup Table 3). Analysis of the *BgtE-20069a* sequence in these isolates revealed five isolates with the reference *BgtE-20069a* sequence as either a single copy gene (three isolates) or as a duplication (two isolates) as well as seven isolates with the R75K gain-of-virulence mutation (Sup Table 3). Phenotyping of two representative isolates for each category on Pm3d#1 transgenic wheat, revealed that isolates with two copies of *BgtE-20069a* exhibit an avirulent phenotype, whereas isolates with a single copy of *BgtE-20069a* showed an intermediate phenotype. Importantly, only isolates carrying BgtE-20069_R75K are fully virulent on Pm3d#1 (Fig. 5g). Taken together, our results show that *Pm3d* indeed recognizes two RALPH effectors, the paralogs BgtE-20069b (AvrPm3d_1/AvrPm3e_2) and BgtE-20069a (AvrPm3d_2) and that virulence on *Pm3d* wheat requires the disruption of both AVRs, by gene deletion of *BgtE-20069b* and the simultaneous presence of the R75K mutation in *BgtE-20069a*.

### Pm3d and Pm3e specificity is defined by single amino acids in the LRR domain

Given that both Pm3d and Pm3e recognize the RALPH effector BgtE-20069b (AvrPm3d_1/AvrPm3e_2), we compared the activity of both *Pm3* alleles towards various mRFP-stabilized BgtE-20069b and BgtE-20069a (AvrPm3d_2) variants upon co-expression in *N. benthamiana*. We selected four variants of BgtE-20069b and three variants of BgtE-20069a found in the parental isolates of our four genetic crosses. Pm3d showed the strongest immune response against BgtE-20069b and BgtE-20069b_T86I, an intermediate response to BgtE-20069b_S74T and BgtE-20069b_H96N and a somewhat weaker response against BgtE-20069a and BgtE-20069a_V108G (Fig. 6a, Sup Fig 11). Only BgtE-20069a_R75K completely evaded recognition by Pm3d (Fig. 6a, Sup Fig 11). In contrast, Pm3e only mounted an immune response against BgtE-20069b and BgtE-20069b_T86I but did not recognize other BgtE-20069b or BgtE-20069a variants (Fig. 6a, Sup Fig 11). We therefore conclude that *Pm3d* clearly excels in recognizing the RALPH effector BgtE-20069b and its paralog BgtE-20069a by mounting a stronger and broader immune response (Fig. 6a). In contrast, Pm3e recognizes fewer BgtE-20069b variants and none of the BgtE-20069a variants but has the unique ability to recognize the structurally unrelated BgtE-5754 (AvrPm3e_1) effector.

Thus, Pm3d and Pm3e exclusively recognize BgtE-20069b_S74T and BgtE-20069a, or BgtE-5754 respectively. This provided the opportunity to study the molecular basis of NLR specificity among closely related, allelic NLR variants. Importantly, the Pm3d and Pm3e NLRs and the non-functional Pm3CS differ at only four amino acid positions, all located within the LRR repeats (Fig. 6b-c). Both Pm3d and Pm3e share the W657 polymorphism (R657 in Pm3CS) in the LRR repeat 4. Additionally, Pm3d contains two polymorphisms, namely R1153 (G1153) and R1356 (T1356) in the C-terminal LRR repeats 22 and 28. Pm3e exhibits a single additional polymorphism, V1332 (E1332) in the C-terminal LRR repeat 27 (Fig. 6b-c).

We introduced each polymorphism either alone or in combinations into the Pm3CS backbone, resulting in 16 chimeric constructs (Pm3_Ch1-16), covering the three natural alleles (Pm3CS, Pm3d and Pm3e) and 13 artificial chimeras (Fig. 6b). We verified protein stability of all chimeras and the absence of autoactivity upon expression in *N. benthamiana* (Sup Fig. 12,13). Next, we co-expressed the 16 NLR variants with BgtE-20069b_S74T and BgtE-20069a (recognized only by Pm3d), and BgtE-5754 (recognized only by Pm3e). Additionally, we tested the 16 chimeras against BgtE-20069b_T86I, which induces strong HR with Pm3d and weaker HR with Pm3e (Fig. 6a).

This analysis revealed that the W657 polymorphism is necessary for the recognition of the RALPH effector BgtE-20069b and its variants. While W657 alone is sufficient to induce a HR response against BgtE-20069b_T86I, it must be combined with other polymorphisms to strongly recognize BgtE-20069b_T86I or to mount an HR response against the more immune-evasive effector variants BgtE-20069b_S74T and BgtE-20069a. In particular, R1356 (found in Pm3d) acts strongly synergistically with W657. The combination of the three amino acids present in Pm3d appears to be optimal as none of the other chimeras recognized the BgtE-20069a variant (Fig. 6b, Sup Fig 14a-c, for details see Sup Note S6).

In contrast, the V1332 polymorphism, naturally present in Pm3e, is crucial for recognizing the effector BgtE-5754, as none of the chimeras lacking it showed an HR response. Although V1332 alone is sufficient, its effect is influenced by other SNPs: the R1356 polymorphism, present in Pm3d but not in Pm3e, acts antagonistically with V1332 in BgtE-5754 recognition, an effect that can be compensated by the simultaneous presence of W657 (Fig. 6b, Sup Fig 14a-c, for details see Sup Note S6).

Taken together, our findings show that the unique abilities of Pm3d and Pm3e rely on only a few polymorphisms in the LRR domain of the NLR protein. Consequentially, the non-functional Pm3CS can be rendered active against BgtE-20069b or BgtE-5754 by the introduction of single polymorphisms, W657 and V1332, respectively. Furthermore, we show that additional polymorphisms can act either synergistically or antagonistically, and that the final combination balances these effects to generate a functional NLR (Fig. 6b-d).

### *Pm3e* and *Pm3d* specificities can be combined in chimeric NLRs

Stacking of *Pm3* alleles using transgenic methods has shown great promise to enhance *Bgt* resistance in wheat^36^. However, several *Pm3* alleles were found to be incompatible due to interallelic suppression effects^37^. We therefore tested for interallelic suppression between *Pm3d* and *Pm3e* in wheat and *N. benthamiana*. In both experimental systems, we observed suppression of *Pm3e* activity by *Pm3d* and vice versa and concluded that combining the two *R* genes in the same wheat line would be of limited use (Fig. 7a,b, Sup Note S7).

Instead, we aimed at combining *Pm3d* and *Pm3e* specificity in a single Pm3 NLR. As a basis, we used the identified *Pm3d/e* chimeras Pm3_Ch12 and Pm3_Ch16 that exhibited *Pm3d-* and *Pm3e*-like specificity (Fig. 6b, Sup Fig. 14). Both chimeras showed an HR response against BgtE-5754 (AvrPm3e_1) which was comparable to Pm3e, suggesting that the chimeras fully incorporate *Pm3e* activity. Furthermore, both chimeras mounted an immune response towards BgtE-20069b_S74T (AvrPm3d_1 variant) which is exclusively recognized by Pm3d. However, the observed HR response was significantly weaker than for Pm3d, in particular for Pm3_Ch12. In addition, both chimeras failed to recognize the BgtE-20069a (AvrPm3d_2) effector (Fig. 6b, Sup Fig. 14). This indicates that Pm3_Ch12 and Pm3_Ch16, although combining *Pm3d-* and *Pm3e*-like specificities do not fully incorporate the Pm3d activity (Fig. 6b-d).

Previously, Stirnweis and colleagues identified two amino acid polymorphisms in the NB-ARC domain (L456P, Y458H) that enhance HR output of Pm3 allelic variants and occasionally increase their recognition spectrum without causing NLR autoactivity^38^. To test if the weak or absent HR response of Pm3_Ch12 and Pm3_Ch16 towards BgtE-20069b_S74T and BgtE-20069a could be improved, we introduced the enhancing polymorphisms L456P and Y458H into the NB-ARC domain of Pm3_Ch12 and Pm3_Ch16, resulting in “enhanced” chimeras enhPm3_Ch12 and enhPm3_Ch16. We subsequently confirmed efficient protein expression and the absence of NLR autoactivity of the engineered NLR constructs (Sup Fig. 15,16).

When tested for recognition of BgtE-5754 (AvrPm3e_1), both enhanced chimeras mounted an HR response comparable to Pm3e and the original Pm3_Ch12 and Pm3_Ch16 chimeras (Fig. 7d). This suggests that recognition of BgtE-5754 is not drastically affected by the two polymorphisms in the NB-ARC. In contrast, both enhanced chimeras performed significantly better against variants of the RALPH AVR effectors BgtE-20069b and BgtE-20069a (Fig. 7e-g). Importantly, the weak HR response of Pm3_Ch12 and Pm3_Ch16 against BgtE-20069b_S74T was significantly increased in the enhanced chimeras, now matching the HR levels mounted by Pm3d (Fig. 7f). Furthermore, the improved chimeras, like Pm3d, also mounted an HR response towards BgtE-20069a (AvrPm3d_2) which evaded recognition by the original chimeras Pm3_Ch12 and Pm3_Ch16 (Fig. 7g).

Next, we tested whether the combined Pm3d- and Pm3e-activity of the enhPm3_Ch12 and enhPm3_Ch16 chimeras can also be observed in wheat under natural *AVR* expression levels. We used particle bombardment to deliver *Pm3* constructs into the epidermis of the susceptible wheat cultivar ‘Bobwhite’ and assessed the haustorium formation index of diagnostic *Bgt* isolates that distinguish *Pm3d* and *Pm3e* resistance, namely *Pm3d*-avirulent/*Pm3e*-virulent isolate CHN_17-40 and *Pm3d*-virulent/*Pm3e*-avirulent isolate CHE_94202 (Fig. 7a-b). As expected, *Pm3d* significantly reduced the haustorium formation rate of isolate CHN_17-40, while it was unaffected by *Pm3e* (Fig. 7h). Congruently, *Pm3e* expression significantly reduced haustorium formation of isolate CHE_94202 while *Pm3d* did not (Fig. 7i). Importantly, both enhanced chimeras enhPm3_Ch12 and enhPm3_Ch16 reduced haustorium formation against both isolates, demonstrating that they are functional in wheat and indeed combine *Pm3d* and *Pm3e* resistance activity (Fig. 7h-i).

These results show that combining Pm3d- and Pm3e-specific polymorphisms in LRR domain with NLR enhancing polymorphisms in the NB-ARC domain allows to incorporate the broad-spectrum resistance activities of Pm3d and Pm3e in a single NLR.

## Discussion

Identifying broad-spectrum resistance genes against pathogens remains an important objective of resistance breeding. We identified *Pm3d* and *Pm3e* as broadly active resistance alleles against globally occurring *Bgt*. Our approach, using transgenic lines expressing individual *Pm3* alleles, allowed the evaluation of resistance without confounding effects that might stem from non-uniform wheat backgrounds. The broad activity of *Pm3d* and *Pm3e* is surprising, given that they are nearly identical to the non-functional *Pm3CS*. Importantly, this broad activity of *Pm3d* and *Pm3e* does not result from constitutive resistance caused by pleiotropic effects due to transgene overexpression, as we identified fully virulent isolates for both alleles. We show that both Pm3d and Pm3e NLRs recognize two *Bgt* effector proteins, which likely contributes to their broad resistance spectrum, as successful resistance evasion by *Bgt* requires simultaneous modifications in both effectors. Indeed, we found that gain-of-virulence mutations in all three avirulence genes exist in *Bgt*, but they do not readily co-occur in the same isolates, thereby forming the basis of the broad activity of *Pm3d* and *Pm3e* worldwide. Nevertheless, the combination of several virulence mutations through sexual recombination in the *Bgt* pathogen might occur upon increased deployment of the *Pm3d* and *Pm3e* resistance genes.

The ability of Pm3d and Pm3e to recognize multiple effectors aligns with growing evidence that NLR/AVR interactions form complex networks that might involve several host and pathogen components^39–41^. For instance, the NLR pair RGA5/RGA4 recognizes two structurally related MAX AVRs (Avr-Pia and Avr-CO39) from *Magnaporthe oryzae*^42,43^. Furthermore, genetic evidence suggests that certain NLRs can also recognize structurally unrelated effectors. For instance, several *Mla* alleles in barley provide resistance against powdery mildew, wheat blast, and stem rust through recognition of, in part, still unidentified effectors^44,45^. Given the divergent effector repertoires of these fungi, the recognized effectors are likely to be structurally unrelated. Therefore, our discovery that Pm3e can recognize two structurally unrelated effectors is of particular significance, as it shows that dual recognition specificity against structurally unrelated fungal effectors can be encoded within a single NLR protein.

The two AvrPm3e effectors belong to two different, structurally defined effector superfamilies. The RALPH superfamily (AvrPm3e_2) contains many characterized avirulence and virulence effectors from *Bgt* and *Bh*^18^. In contrast, the AvrPm3e_1-like superfamily, first described based on sequence similarity over a decade ago^46^ remains functionally uncharacterized. This is surprising given its ∼13% contribution to the *Bgt* effectome. An independent study by Seong et al. identified a structural effector superfamily in *Bh*, termed „cluster 40“, that corresponds to our definition of AvrPm3e_1-like effectors^34^. Of note, the authors did not find evidence for the existence of this superfamily in other phytopathogenic fungi, therefore supporting our findings that the AvrPm3e_1-like effectors are an innovation of the *Blumeria* lineage.

Recently, 9o9, an AvrPm3e_1-like effector of *Bh*, was identified as an interaction partner of the barley *Bh*-susceptibility factor RACB, a small monomeric GTPase^47^. Reminiscent of our finding for BgtE-5754 (AvrPm3e_1), the authors failed to predict a clear signal peptide for the 9o9 effector and overexpression of the full-length 9o9 effector increased the penetration rate of *Bh* in barley. Similarly, the full-length BgtE-5754, but not any of the tested N-terminal truncation variants corresponding to putative signal peptide prediction, triggered a Pm3e-mediated HR in *N. benthamiana*. This suggests that the N-terminal region is indeed part of the mature AvrPm3e_1 protein and might not function as a classical signal peptide. Consequently, the mechanism by which effectors such as BgtE-5754 or 9o9 exit the fungal cell and reach the host cytoplasm remains unclear^48^. Given increasing evidence from other phytopathogens for alternative secretion pathways, it is likely that similar mechanisms also exist in *Blumeria*^49,50^. The identification of both an avirulence and a virulence factor among the AvrPm3e_1-like effectors suggests that, like RALPH effectors, they represent important components in the effectome of *Bgt* and *Bh*. Therefore, it is likely that the *AvrPm3e_1*-like superfamily contains additional AVRs that have remained undetected, for instance, because previously performed effector screens have focused on the RALPH effectors exclusively^11^.

We found that the specificity of Pm3d and Pm3e towards BgtE-5754 and BgtE-20069a/b effectors, are encoded withing single amino acid changes compared to the completely inactive Pm3CS. Similarly, in the NLR Sr50, a single substitution was necessary and sufficient to restore recognition towards a non-recognized AvrSr50 variant. Reminiscent to our discovery that all other polymorphisms modulate rather than define the HR response, additional mutation in Sr50 also acted synergistically on HR amplitude^51^. The fact that specificity towards structurally unrelated effectors can be encoded by single amino acid changes in an otherwise inactive NLR backbone such as Pm3CS, indicates that recognition of additional structural folds, both from *Bgt* and beyond, might be engineerable into existing NLR receptors. In addition, it suggests that NLRs are more versatile than previously assumed and that other NLRs might also contain the intrinsic capacity to recognize structurally unrelated effectors.

Finally, we engineered chimeric NLRs with the dual Pm3d/Pm3e specificity through a combination of natural polymorphisms, thereby overcoming interallelic suppression effects observed between the two Pm3 alleles. These findings highlight the potential of NLR engineering when pyramiding of natural *R* genes is hampered by antagonistic effects.

Furthermore, the successful combination of naturally occurring polymorphisms in the NB-ARC and LRR domain resulting in our enhanced Pm3 chimeras (enhPm3_Ch12 and enhPm3_Ch16), exemplifies how allelic diversity can inform NLR engineering to alter recognition specificity or modulate the HR output. Such approaches, informed by naturally occurring diversity within allelic series, might be particularly useful for the engineering of NLRs where high-resolution structural data is missing, as is the case for the majority of NLR receptors^52,53^. In addition, the observation that only five or six polymorphisms are sufficient to render the non-functional Pm3CS NLR into the highly functional enhPm3_Ch12 or enhPm3_Ch16 chimeras with the ability to recognize three AVR effectors, opens the possibility for direct engineering of these new extended specificities in elite breeding material. This could for instance be achieved via targeted base editing of the naturally occurring *Pm3CS* allele that is found in many wheat breeding lines^25,26,54^. Previous studies have reported that NLR engineering efforts based on HR output measured in *N. benthamiana* did not consistently translate into resistance in the corresponding homologous host^55,56^. Therefore, our finding that the chimeric receptors are functional in wheat and restrict pathogen penetration is reassuring and renders them promising candidates for stable expression in wheat.

In summary, our study illustrates that allelic series are a valuable source of diversity that can be exploited in disease resistance breeding. First, allelic series may contain so far undiscovered broadly active alleles that can be directly incorporated into breeding programs. Secondly, polymorphisms within allelic series provide valuable information to guide NLR engineering. For both approaches, highly diverse allelic series rich in unexplored diversity, such as the *Pm3* locus, are particularly well suited. Furthermore, the ability of Pm3 NLR to recognize structurally unrelated effector folds, combined with their amenability to NLR engineering, makes them particularly valuable for breeding. We propose that the identification of naturally occurring, or the creation of engineered NLRs with multiple recognition specificities represent a promising strategy for resistance breeding. Such NLRs might act similarly as *R* gene stacks by reducing the likelihood of resistance breakdown, thereby contributing to more durable wheat protection.

## Material and Methods

### Plant material

All transgenic wheat lines used in this study have been previously generated and express individual *Pm3* alleles under a maize ubiquitin promoter in the susceptible wheat cultivar ‘Bobwhite SH 98 26’. These transgenic lines (alongside respective sister lines) were selected as best performing lines exhibiting high transgene expression and race-specific *Bgt* resistance in the absence of pleiotropic effects^28–31,36^. Specifically, the transgenic lines Pm3a#1, Pm3b#1, Pm3d#1, Pm3e#2, Pm3g#1 and Pm3CS#19 were used in this study. For Pm3d#1 and Pm3e#2 the corresponding sister lines Sd#1 and Se#2 were included as controls.

To generate Pm3d#1 x Pm3e#2 F1 hybrids, homozygous Pm3d#1 and Pm3e#2 parental lines were crossed and the resulting F1 progeny were phenotypically assessed for *Pm3d*- and *Pm3e*-specific resistance.

### Fungal isolates and virulence phenotyping

All *Bgt* isolates used in this study represent twice single spore isolated, genetically uniform cultures that have been described in previous studies^15,16,27,32^. Isolates were maintained on detached leaves of the susceptible wheat cultivar ‘Kanzler’ placed on 0.5% agar plates (PanReac AppliChem), supplied with 4.23 mM benzimidazole^57^.

Virulence phenotypes were assessed on detached leaf segments cut from primary leaves of 10 day old seedlings that were placed on 0.5% agar plates before infection (0.5% food grade agar (PanReac AppliChem), 4.23 mM benzimidazole) and incubated at 20°C^57^. Virulence scoring was performed 8-11 days post infection, depending on the growth speed of individual *Bgt* isolates on the susceptible wheat cultivar ‘Kanzler’, serving as an infection control. Virulence phenotypes reported or depicted in Fig. 1b, Fig. 2, Fig. 5a/b/g were determined on three to six biological replicates based on the observed mildew leaf coverage (MLC) and classified in five qualitative categories: 1 (100% MLC), 0.75 (75% MLC), 0.5 (50% MLC), 0.25 (25% MLC) and 0 (0% MLC). Phenotyping results are summarized in Supplementary Dataset S1. To assess the more quantitative interallelic suppression phenotype in the Pm3d#1 x Pm3e#2 F1 hybrids, 18 biological replicates were assessed and individually scored based on observed MLC in 11 categories ranging from 0% to 100%. Representative images were chosen for the depiction of virulence phenotypes throughout the manuscript.

### Avirulence depletion assay

Avirulence depletion mapping was performed as previously described^16^. In brief, the AD assay relies on the generation of a haploid F1 progeny population by crossing two *Bgt* isolates with opposite virulence phenotypes on the *R* gene of interest, followed by selection of the F1 population on a resistant host cultivar thereby resulting in depletion of F1 progenies carrying the parental avirulent genotypes. AVR-depleted F1 populations are then bulk sequenced and compared to a non-selected control bulk to identify genomic regions with strong deviations from a 1:1 segregation of parental markers, indicating the presence of an *AVR* factor.

Sexual crosses were performed between the isolates CHN_52-27 (MAT1-1) x CHN_17-40 (MAT1-2), IRN_GOR_2 (MAT1-1) x CHN_17-40 (MAT1-2) and CHE_94202 (MAT1-1) x USA_2 (MAT1-2) by co-infection of the susceptible wheat cultivar ‘Kanzler’ in a greenhouse chamber^58^. The cross between CHN_52-27 (MAT1-1) and CHVD_042201 (MAT1-2) was performed for a previous study following the same protocol^16^. Arising chasmothecia were harvested, dried and ascospores ejected as previously described^16^. Arising F1 progeny were then subjected to either selection on Pm3e#2 or Pm3d#1 transgenic lines (*Pm3*-selected bulks) or grown on ‘Kanzler’ (non-selected bulk). Conidiospores of individual bulks were harvested 10 days post infection and fungal DNA was extracted using a previously described CTAB/phenol chloroform extraction protocol^9^. Bulk DNA was sequenced on a NovaSeq X Plus platform with our commercial partner Novogene (https://www.novogene.com) at a coverage of approximately 120x.

To perform the AD bioinformatics pipeline, Illumina reads of the parental isolates of each cross were aligned to the reference genome of the respective avirulent parent of each cross and high-quality markers defined. The respective genome assemblies have been described previously^59^. Mapping of Illumina data, SNP calling and high-quality marker definition were performed as described previously^16^ using bowtie2 v2.4.4. and FreeBayes v1.3.6. The corresponding Python script is available from: https://github.com/MarionCMueller/AD-assay/blob/main/scripts/Script_BSA_SNPcall.v8.py. Following marker definition, sequencing data of *Pm3*-selected and non-selected bulks were mapped to the respective avirulent parental genome assembly, followed by determination of parental genotype ratios. To test for deviations from the expected 1:1 ratio indicating avirulence depletion, a G-test of goodness-of-fit was performed using the R package RVAideMemoire and the G.test() command as described in^16^. The full code is available under https://github.com/MarionCMueller/AD-assay/blob/main/scripts/BSA_Blumeria_functions.R

To define genomic intervals of *AVR* loci, the percentage of alternative alleles over five consecutive polymorphic sites were averaged and the largest interval that surpassed 90%, 95%, and 98% of reads, respectively, was extracted. For each cross, the interval with the highest selection was retained. To define the overlapping interval between all crosses, 400 bp of the flanking regions of each interval were extracted and subsequently aligned against the reference genome Bgt_genome_v3_16^33^ using blastn v2.12.0^60^.

### AvrPm3 haplovariant mining

To assess haplovariant diversity of individual AvrPm3 effectors, Illumina resequencing datasets were mapped to the *Bgt* reference assembly of CHE_96224 as previously described^27^. For each isolate, mapped reads were visualized using integrative genomics viewer IGV (v2.8.6)^61^ and haplovariants defined manually. For isolates with ambiguous results due to gene duplications or sequence insertions (i.e. 9 bp insertion in AvrPm3e_1), determined haplovariants were verified by PCR amplification and subsequent in-house Sanger sequencing, using the primers listed in Sup Table 4.

### Gene cloning

To produce Gateway compatible entry clones of individual effectors and effector variants, their coding sequence was codon-optimized for *N. benthamiana* with the codon-optimization tool of Integrated DNA technologies (https://eu.idtdna.com/pages/tools/codon-optimization-tool) and subsequently produced with flanking attL1/attL2 sites by gene-synthesis (commercial partners: BioCat GmbH, Thermo Fisher Scientific, Integrated DNA technologies), PCR-based site directed mutagenesis (single polymorphisms) and/or in-Fusion cloning (mRFP fusions). BgtE-20069a and BgtE-20069b variants were cloned without their predicted signal peptide (SignalP4.0)^62^, whereas BgtE-5754 variants were cloned with their full-length coding sequence unless indicated otherwise. The DNA sequences of all effector constructs used in this study are compiled in Supplementary Dataset 2. Pm3CS-HA, Pm3d-HA, Pm3e-HA and Pm3g-HA genomic DNA sequences^11^ were cloned into the Gateway compatible pENTR plasmid using in-Fusion cloning (Takara Bio). Subsequently, PCR based site-directed mutagenesis with primers listed in Sup Table 4 was used to introduce individual amino acid polymorphisms resulting in the chimeric constructs Pm3_Ch1-16, enhPm3_Ch12 and enhPm3_Ch16.

All entry clones described above were subsequently mobilized into the binary vector pIPKb004^63^ using Gateway LR clonase II (Invitrogen). Resulting expression constructs were sequence verified by in-house Sanger sequencing (effectors) or full plasmid sequencing (*R* genes) with our commercial partner (Microsynth AG) and transformed into *Agrobacterium tumefaciens* GV3101 using freeze-thaw transformation^64^.

For bombardments of wheat epidermal cells, entry clones containing Pm3CS-HA, Pm3d-HA, Pm3e-HA, enhPm3_Ch12-HA and enhPm3_Ch16-HA (see above) were mobilized into a Gateway-compatible pGY1 expression vector^65^ using Gateway LR Clonase II (Invitrogen).

The expression constructs for Pm3a-HA, Pm3b-HA, Pm3c-HA and Pm3f-HA in the binary vector pIPKb004 have been previously described^9,11^. The RNA silencing inhibitor p19 has been previously described^66^.

### AVR validation and HR quantification

*Agrobacterium*-mediated expression in *N. benthamiana* was achieved as previously described^11^. For *AVR-R* co-expression, *Agrobacteria* at an OD600 of 1.2, carrying effector constructs, *R* gene constructs or the silencing inhibitor p19 were mixed in a 4:1:1 ratio prior to infiltration. The HR response was imaged and quantified at 4 days post infiltration (unless indicated otherwise) using a conventional camera and/or a Fusion FX imaging System (Vilber Lourmat, Eberhardzell, Germany) and subsequently quantified using Fiji^11,67^. All co-expression experiments were performed three times using n=6 biological replicates per assay, resulting in a total of n=18 HR measurements for quantification and statistical analysis. Representative images were chosen to depict the HR phenotypes throughout the manuscript.

### Western blotting

Effective production of effector or NLR proteins in *N. benthamiana* was tested using the *Agrobacterium* infiltration protocol described above. Single constructs were mixed with p19 silencing inhibitor in 4:1 ratio prior to infiltration. Infiltrated plant tissue was harvested at 2 days post infiltration and proteins were extracted in from 4 x 2 leaf discs (0.5cm diameter) in 150 μl 2x Lämmli buffer (62 mM Tris–HCl pHC6.8, 10% glycerol, 2% SDS, 0.02% bromophenol blue, 1% β-mercaptoethanol) and boiled for 5 min at 95°C followed by brief centrifugation (12’000rpm, 10min) to remove cell debris. Protein extracts were separated on either 8% (NLR detection) or 4-20% gradient (effector detection) SDS polyacrylamide gels respectively and subsequently blotted to a nitrocellulose membrane (Amersham Protran 0.2 μm NC). Detection of HA-tagged Pm3 proteins was achieved by using 1:3,000 diluted anti-HA-HRP antibody (rat monoclonal, clone 3F10, Roche). Detection of FLAG-or 3xFLAG-tagged effectors was achieved by using 1:3,000 diluted anti-FLAG-HRP antibody (mouse monoclonal, clone M2, Sigma Aldrich). HRP chemiluminescence was induced with WesternBright ECL HRP substrate (Advansta) and imaged on a Fusion FX imaging System (Vilber Lourmat, Eberhardzell, Germany).

### Bombardment of wheat epidermal cells

The pGY1 plasmids carrying Pm3CS-HA, Pm3d-HA, Pm3e-HA, enhPm3_Chr12-HA and enhPm3_Chr16-HA (see above) were extracted using the NucleoBond Xtra Midi Kit (Macherey-Nagel) according to manufacturer instruction. Particle bombardments were performed as described in^24^. To do so, 1.25 µg of Pm3 plasmids DNA as well as 1.25 µg pUbi-GUS plasmid DNA were attached to gold particles (11 μl of 25 mg/ml gold particles with 1 μm diameter, Bio-Rad, Hercules, USA) and shot onto 9-day-old first leaves of the susceptible wheat cultivar ‘Bobwhite’ using the PDS-1000/HE^TM^ (Bio-Rad, Hercules, USA) particle gun. Per experimental replication, two times five leaf segments were bombarded for each construct and isolates combination. One hour post bombardment leaf segments were infected with conidiospores of *Bgt* isolates CHN_17-40 or CHE_94202. Leaf segments were vacuum infiltrated in GUS-staining solution (0.1 M NaCHPOC/NaHCPOC pH 7.0, 0.01 M EDTA, 0.005 M potassium hexacyanoferrate (II), 0.005 M potassium hexacyanoferrate (III), 0.1% (v/v) Triton X-100, 20% (v/v) methanol, 0.5 mg/mL 5-bromo-4-chloro-3-indoxyl-β-D-glucuronic acid) 44 h post inoculation and subsequently incubated at 37°C for 3–5 h before destaining in 70% ethanol. For microscopic evaluation, leaves were submerged in Calcofluor White solution^47^. Haustorium formation rates were determined in GUS-positive cells attacked by a single *Bgt* spore. For each construct, at least 25 interactions were counted. The experiment was performed three times.

### Bioinformatics analysis

Pfam domains of full-length BgtE-5754 were predicted using the InterPro webserver (https://www.ebi.ac.uk/interpro/)^68^. Signal peptides for protein sequences of BgtE-5754 and BgtE-20069 were predicted using SignalP 4.1, 5.0, 6.0 and DeepTMHMM webservers with default parameters^62,69–71^.

To define effectors with structural similarity to AvrPm3e_1 in *Bgt*, the structure of the 844 candidate *Bgt* effectors^33^ was predicted using AlphaFold2^72^. Subsequently, the best scoring model for each effector was compared pairwise for structural similarity using TM-align (v20220412)^73^. In a first step, hits with a pairwise TM-align score > 0.6 when compared to BgtE-5754 were considered as members of the AvrPm3e_1-like superfamily. To identify additional effectors with structural similarity, iterative searches against all previously defined AvrPm3e_1-like family members were performed (TM-align score > 0.6) until saturation. The resulting AvrPm3e_1-like superfamily contained 112 members belonging to the previously defined effector families E001 (including BgtE-5754), E027, E040 and E061 (nomenclature of^33^). The same procedure was applied using the protein structural model of BgtE-20069b to identify RALPH effectors.

To search for AvrPm3e_1-like effectors in *Bh*, the 112 AvrPm3e_1-like proteins in *Bgt* were compared with previously predicted AlphaFold structures of the *Bh* effectome^34^ using TM-align (score >0.6). The Foldseek server^74^ was used to search for AvrPm3e_1-like effectors in powdery mildew fungi outside the *Blumeria* genus. To do so, BgtE-5754 and randomly selected five additional members of family E001 and two members of family E027, E040, E061 were searched against the AlphaFold/UniProt50_v4 database with search results restricted to Erysiphales. The resulting hits, irrespective of significance score, were retrieved and compared in pairwise similarity searches against the 112 AvrPm3e_1-like effectors in *Bgt* using TM-align (score >0.6). Accession numbers of analysed protein structures can be found in Supplementary Dataset 3.

Signal peptide prediction was performed for all proteins of *Bgt* reference isolate CHE_96224 (Bgt_CDS_v4_23, available at: https://doi.org/10.5281/zenodo.701850175)using SignalP 6.0^70^ in the mode --organism eukaryote --mode slow-sequential with default thresholds for signal peptide definition.

Expression analysis of CHE_96224 effectors was performed using publicly available RNA-seq data from isolate CHE_96224 at two days post inoculation on the susceptible wheat cultivar “Chinese Spring” (SRA accession numbers: SRX7003606-SRX7003608). The coding sequence of CHE_96224 annotation Bgt_CDS_v4_23 (see above) was indexed using the Salmon v1.4.0^76^ *index* command, using the chromosomes as decoys and with the option –keepDuplicates. Subsequently, the transcripts were quantified using *salmon quant* with the - l A option and RPKM values calculated with the edgeR package v3.40.2^77^ using the command rpkm(). Finally, the average of the three biological replicates was calculated.

Structural prediction of AVR proteins was performed using the AlphaFold3 server (https://alphafoldserver.com/)^78^. For BgtE-5754, the full-length protein sequence was used, whereas structural prediction of BgtE-20069b (AvrPm3d_1), BgtE-20002 (AvrPm3b/AvrPm3c), and Bgt_avrF2_7 (AvrPm3a/AvrPm3f) was performed on their respective mature proteins (SignalP 6.0)^70^. Predictions for all proteins were of sufficient quality (pTM score >0.8), except for the BgtE-20002 protein that had a pTM score of 0.33. Inspection of the multiple sequence alignment (MSA) for BgtE-20002 underlying the prediction revealed a low number of proteins in the MSA, likely due to the UniProt database being outdated for *Bgt*. High-quality prediction of the BgtE-20002 structure (pTM score >0.8 was achieved by using the ColabFold_AlphaFold2 pipeline^79^ with a custom MSA generated with the HHblits Toolkit^80,81^, containing all high-quality protein sequences of the BgtE-20002 effector family E018. All resulting structural predictions were visualised with ChimeraX v1.8^82^.

## Acknowledgments

We thank Prof. Dr. Ralph Hückelhoven for the support of the work at the Technical University of Munich and for discussions of results. We thank Esther Jung at the University of Zurich and Veronika Schmitt at the Technical University of Munich for technical assistance. We are also grateful to Johannes Graf for support with bioinformatic analyses. Furthermore, we acknowledge Karl Huwiler and Urs Somalvico for their assistance with plant growth and maintenance.

## Author contributions

L.K., M.C.M. and B.K. designed and conceived the study. L.K., J. I. and M.C.M performed the experiments. M.C.M, L.K., and M.H. performed bioinformatic analyses. Z.B., A.G.S., U.S., J.J., and F.M. contributed biological resources and sequencing datasets. B.K., L.K., M.C.M., and T.W. supervised work. L.K., M.C.M and B.K. wrote the manuscript. All authors read and approved the final version of manuscript.

## Data availability

Sequencing reads produced in this study for the avirulence depletion assay are deposited at the sequence read archive under accession number PRJNA1125842. Sequencing reads of the mutant Bgt_FAB are available under accession number PRJNA1016363. Predicted protein structures of *Bgt* effectors are available at Zenodo (https://doi.org/10.5281/zenodo.17276912)

