## Supplementary Material for "Dual recognition of structurally unrelated mildew effectors underlies the broad-spectrum resistance of Pm3e in wheat"

[2] Current address: School of Molecular Biosciences, University of Glasgow, Glasgow, UK

[3] Current address: Centre for Crop Health, University of Southern Queensland, Australia

**Supplementary Note S1**

The avirulence depletion assay relies on depleting a haploid F1 population of individuals carrying the avirulent allele upon selection on a resistant host cultivar followed by identification of AVR associated genomic regions indicated by deviations from a 1:1 segregation of parental markers. To ensure that observed depletion signals are indeed specific to the *R*-gene mediated selection pressure and hence indicate the genomic location of sought-after *AVR* genes, the AD assay relies on an unselected control bulk generated by growing the same F1 population on a susceptible cultivar devoid of *R* genes.
In the four genetic crosses performed in this study to identify *AvrPm3e*-components, two F1 populations, namely CHN_52-27 x CHN_17-40 and CHN_52-27 x CHVD_042201, did not show any signal of allele depletion on the control bulk on susceptible cultivar "Kanzler" (Figure 2). However, the non-selected bulks of the remaining two crosses (IRN_GOR2 x CHN_17-40 and USA_2 x CHE_94202) exhibited depletion signals in one region on Chr-06 (IRN_GOR2 x CHN_17-40) or on chr_07 (USA_2 x CHE_94202) respectively. These findings indicate that these two crosses segregate for additional genetic factors affecting pathogen growth that are not associated with virulence on *Pm3e,* and we consequently excluded those loci from the analysis aiming at the identification of *AvrPm3e* components.
One possible explanation of *R*-gene independent allele depletion in F1 bulks is differential strain aggressiveness (i.e. presence of virulence factors) between the parental isolates. Progenies with increased aggressiveness would outcompete the less fit progenies even on the control line 'Kanzler' and therefore cause the deviation from the 1:1 ratio in regions underlying this differential aggressiveness. Since in our assay, F1 progenies undergo only two rounds of sexual reproduction before sequencing, it is unlikely that such a quantitative difference in strain aggressiveness would lead to complete fixation of the stronger virulence allele. Indeed, for the *Pm3e*-independent loci on Chr-06 in the IRN_GOR2 x CHN_17-40 (Figure 2), we observed an overrepresentation of CHN_17-40 parental makers in the locus to a prevalence of approximately 80%, Therefore, this locus likely contains a virulence factor in CHN_17-40.
In contrast, the depletion signal observed on Chr-07 in USA_2 x CHE_94202 was near complete with close to 100% of obtained reads carrying the USA_2 genotype in this genomic region. We consider it unlikely that a quantitative virulence factor would result in full fixation of the advantageous trait within two rounds of asexual propagation. We initially suspected a technical artifact as the cause of the observed phenomenon. Specifically, we considered that our genetic marker definitions, based on Illumina reads from parental isolates, might not fully reflect the genotype of isolates used for crossing. We hypothesized that CHE_94202 may have acquired a deletion during repeated asexual propagation, leading to the absence of these markers in the F1 population. However, the presence of the Chr-07 locus in both parental isolates was confirmed by Sanger sequencing, demonstrating the accuracy of the Illumina data and ruling out such a technical artifact.
Therefore, we considered biological phenomena that could lead to such a strong allele depletion. One possibility is that progenies carrying the CHE_94202 genotype fail to complete ascospore development or maturation due to a disrupted gene essential in these processes and located within the region on Chr-07. Alternatively, the region on Chr-07 may harbour a spore killer mechanism, as observed in other ascomycete fungi[1], whereby spores with the USA_2 genotype

actively eliminate those with the CHE_94202 genotype during development, resulting in complete fixation of USA_2 markers at this locus.
Our observations highlight the importance of generating a non-selected control bulk as part of the AD assay pipeline to detect allele depletions that are independent of *R* gene-mediated selection and therefore should be excluded from downstream analyses focusing on AVR identification.

### Supplementary Note S2

Our attempts to predict a signal peptide in BgtE-5754 yielded varying results depending on the software used. For instance, SignalP 6.0 predicted a signal peptide with cleavage between amino acids 16 and 17, however, with low probability (0.574). The same cleavage side was predicted by SignalP 4.1, however, the prediction only slightly surpasses the threshold for SP detection. In contrast, the SignalP 5.0 predicted the cleavage site after aa 15, albeit with very low probability (0.17). Finally, DeepTHMM predicted a SP of size 13 (Sup Fig 1a). This is in contrast to the consistent prediction of cleavage of a signal peptide between amino acids 21 and 22 in the RALPH effector BgtE-20069b (AvrPm3d) (Sup Fig 1a).
We tested truncated versions of BgtE-5754-A1 and A2, lacking the amino-terminal 13 or 16 residues, for their ability to trigger Pm3e-dependent HR. In contrast to their full-length versions, none of these truncations triggered an HR response, thus indicating that only full-length BgtE-5754 proteins are recognized by Pm3e (Sup Fig 1b). This furthermore suggests that the N-terminal part of BgtE-5754 does not constitute a signal peptide but rather a part of the mature BgtE-5754 protein.
Interestingly, an independent study in the closely related *Bh*, identified a structurally related effector protein that did not display a clear signal peptide[2]. This effector, termed 9o9, was found in a screen for interaction partners of the barley susceptibility factor RACB, a Rho-of-plants small GTPAse, and shown to contribute to *Bh* infection success if expressed as a full-length protein in barley cells. Importantly, both RACB and Pm3 NLRs were shown to be localized intracellularly[2,3] suggesting that 9o9 and BgtE-5754 effectors reach the plant cytoplasm despite the lack of a clear signal peptide. While little is known about unconventional secretion pathways in *Blumeria*, the growing evidence for effector secretion mechanistically distinct from the classical secretory pathway in other phytopathogenic fungi[4], suggests that such pathways may also exist in *Blumeria*.

### Supplementary Note S3

To verify that virulent BgtE-5754 variants are effectively produced upon *Agrobacterium*-mediated expression in *N. benthamiana*, we performed epitope tagging followed by western blot analysis of four BgtE-5754 variants (A1, A2, V1 and V2). To do so, the full-length BgtE-5754 C-terminally was fused with mRFP-3xFLAG. All four variants produced equal amounts of protein (Sup Fig 3), indicating that the underlying amino acid polymorphisms in the virulent variants BgtE-5754-V1 and V2 do not negatively affect protein production but are involved in Pm3e evasion. Our western blot analysis furthermore revealed that all four tested BgtE-5754 variants are prone to posttranslational processing or protein degradation in *N. benthamiana*, with a large band corresponding to the predicted BgtE-5754-mRFP-3xFLAG protein size of ~70 kDa and up to three smaller fragments (Sup Fig 3). It is likely that such truncated products are caused by lack of protein stability in *N. benthamiana* or by posttranslational processing, potentially related to the absence of a clear signal peptide in BgtE-5754 (see Sup Note S2).

**Supplementary Note S4**

Within the six parental isolates used for mapping of the *AvrPm3e_1* component, we found three avirulent variants (BgtE-5754-A1/A2/A3) and three virulent variants (BgtE-5754-V1/V2/V3), indicating that the gene is diverse within the *Bgt* population. Screening the 220 global isolates described by Sotiropoulos and colleagues[5] identified four additional variants with single amino acid polymorphisms and one with a small insertion (Fig. 3e, Sup Dataset 1). Three variants defined by the polymorphisms K105N/R256Q, K251R/R256Q/Q335H/E367G, and a three amino acid residue insertion (RLD) at position 18 in addition to R256Q, triggered a *Pm3e*-dependent HR response and were consequently designated BgtE-5754-A4, A5 and A6 respectively. In contrast, the variants with polymorphisms R256Q/R331Q and R256Q/T265K/H299R/R309I were not recognized by Pm3e and hence named BgtE-5754-V4 and V5, respectively (Fig. 3e). In summary, this analysis demonstrates that virulent variants of BgtE-5754 (AvrPm3e_1) are present in the global *Bgt* population.
Next, we investigated which amino acid polymorphisms naturally present in BgtE-5754 are responsible for evasion of Pm3e-mediated HR in *N. benthamiana*. To do so, we introduced all single amino acid substitutions found in the *Bgt* diversity panel into the BgtE-5754-A1 variant and tested the resulting 14 constructs for induction of a Pm3e-mediated HR response in *N. benthamiana*. Nine of these 14 constructs did not affect the Pm3e-mediated HR (Fig. 3f, Sup Fig 4). Interestingly, five of the mutations (H299R, R309K, R309I, R312M, and Q335H), all located within a short C-terminal stretch of the BgtE-5754 protein, completely abolished Pm3e recognition (Fig. 3f, Sup Fig 4). Because there is no experimental structural data for BgtE-5754 (AvrPm3e_1), we predicted its structure with Alphafold3 and investigated the location of the immune-disruptive amino acids (Fig. 3f). Interestingly, four of the immune-evasive amino acid polymorphisms affect three positions within the same predicted alpha helix in the predicted BgtE-5754 structure with the fifth polymorphisms located in the adjacent alpha helix (Fig. 3f). The observation that all immune evasive polymorphisms are located in close proximity suggests that this C-terminal domain is involved in BgtE-5754 recognition.

**Supplementary Note S5**

Since BgtE-5754 (AvrPm3e_1) is significantly larger than all previous avirulence effectors recognized by the Pm3 NLRs, we characterized this effector in more detail. First, we examined whether other effectors in *Bgt* share a similar structural fold. To do so, we predicted the structure of all 844 *Bgt* candidate effectors[6] using AlphaFold2, excluding the signal peptides prior to prediction. Using TM-align and the BgtE-5754 structural model as a reference, we identified 79 candidate effectors with pairwise structural similarities (TM-align score > 0.6). The structurally related effectors belong to two different *Bgt* candidate effector families: E001 (harboring BgtE-5754) and E027 (nomenclature of Müller et al. (2019)[6]). Next, we performed iterative structural similarity searches using all previously identified effector models. This process was repeated until no additional structurally similar effectors were found. After four iterations, the final set of structurally similar effectors comprised 112 proteins, belonging to four candidate effector families: E001, E027, E040, and E061. We will subsequently refer to this group of effectors as the AvrPm3e_1-like superfamily. To compare this superfamily to the well-described RALPH effectors, we repeated the iterative similarity searches with the BgtE-20069b protein, previously identified as AvrPm3d[7], resulting in a structurally defined group of 321 canonical RALPH effectors. We subsequently compared the two effector gene superfamilies with regard to expression levels, gene models, protein size/structure and the presence of a signal peptide.

Expression analysis based on a previously generated RNA-seq dataset from the reference isolate CHE_96224 two days post infection, revealed that RALPH effector genes are, on average, expressed at much higher levels compared to genes in the AvrPm3e_1-like superfamily (Fig 3g). Furthermore, while RALPH effector genes were previously shown to contain a single highly conserved intron[8], the majority of AvrPm3e_1-like effectors are encoded by a single exon (Fig 3h). On the protein level, the small RNase-like structure of RALPH effectors consists of an average of 150 amino acid residues. In contrast, AvrPm3e_1-like effectors are significantly larger, averaging 331 amino acids and exhibiting no structural similarities with RNases (Fig. 3f, i).

Lastly, signal peptide prediction on members of both superfamilies revealed that, while RALPH effectors predominantly contain a well-defined signal peptide, this feature is absent (or cannot reliably be detected) in 67% of the AvrPm3e_1-like superfamily, including BgtE-5754 (AvrPm3e_1) itself (Fig 3j, Sup Fig 1). Taken together these findings indicate that BgtE-5754 belongs to a highly expanded superfamily in *Bgt* which is fundamentally distinct from all previously identified RALPH AVR effectors.

Next, we studied whether AvrPm3e_1-like effectors are also found in the *Blumeria* sister species *Blumeria hordei* (*Bh*) (barley powdery mildew), and potentially in other powdery mildew fungi. We analyzed 993 predicted *Bh* candidate secreted protein structures generated in a previous study[9]. Using the TM-align pairwise structural similarity approach (TM-align score > 0.6), we identified 82 *Bh* proteins with similar predicted folds, indicating they belong to the AvrPm3e_1-like superfamily. Interestingly, Seong et al. previously classified 87 Bh proteins into a structural cluster (cluster_40), which was not found in other phytopathogenic fungi[9]. Importantly, all 82 AvrPm3e_1-like *Bgt* proteins identified in our analysis fall within the *Bh* cluster_40, indicating that the two approaches identified the same structural fold independently. While the study of Seong & Krasileva did not find evidence for the existence of structurally similar proteins in other phytopathogenic fungi, their diversity panel contained a single powdery mildew species (*Bh*). In a next step, we therefore aimed at exploring whether powdery mildew species outside the *Blumeria* genus contain AvrPm3e_1-like effector proteins. We searched the proteomes of all deposited fungal species in the order of *Erysiphales* using the Foldseek algorithm[10] for similarities with the BgtE-5754 model and 11 additional randomly selected models of AvrPm3e_1-like proteins (five from family E001, two from family E027, E040 and E061 each) (see Supplementary Dataset 3). In the first step, we did not apply any specific filtering steps but retrieved all non-*Blumeria* protein models identified by Foldseek. With this strategy, we retrieved 21 protein models. We then performed local TM-align searches against the 112 AvrPm3e_1-like effectors from *Bgt*. We found that none of the 21 sequences showed a pairwise TM-align score above 0.6. We therefore concluded that the AvrPm3e_1-like superfamily is expanded specifically in the *Blumeria* genus.

**Supplementary Note S6**

To test which of the four amino acid differences between the active Pm3d and Pm3e NLRs and the inactive Pm3CS mediate specificity towards the different AvrPm3d/e effectors, we created 16 chimeric NLRs (Pm3_Ch1-16, including the natural alleles Pm3CS, Pm3d and Pm3e), each incorporating a distinct combination of these amino acid changes. We then assessed the ability of these NLRs to recognize BgtE-5754-A1 (recognized exclusively by Pm3e) and the three RALPH effector variants BgtE-20069b_T86I (recognized by Pm3d and Pm3e), BgtE-20069b_S74T, and BgtE-20069a (both recognized exclusively by Pm3d) in *N. benthamiana* (Figure 6b-d, Sup Fig 14). This analysis revealed that the ability of Pm3d and Pm3e to recognize BgtE-20069b_T86I (AvrPm3d_1/AvrPm3e_2) depends largely on the presence of W657, which is shared by both alleles. None of the chimeric NLRs carrying an R at position 657 such as Pm3CS, were able to mount an immune response against BgtE-20069b_T86I, whereas the introduction of W657 alone into the Pm3CS backbone (Pm3_Ch2) was sufficient to induce an HR response, albeit somewhat weaker than with the Pm3e and particularly the Pm3d NLRs. None of the other polymorphisms, when introduced alone into Pm3CS were sufficient to generate a functional NLR allele against BgtE-20069b_T86I (Pm3_Ch3, Pm3_Ch4, Pm3_Ch5). The weak HR response of Pm3_Ch2 (W657 alone) indicated that other polymorphisms found in Pm3d and Pm3e contribute synergistically to increase NLR activity against BgtE_20069b_T86I. Indeed, all other polymorphisms, when combined with W657 (Pm3_Ch6, Pm3_Ch7 (=Pm3e), Pm3_Ch8), resulted in an improved HR response (Fig. 6b-d, Sup Fig 14). In particular, the combination of W657 and R1356 (Pm3_Ch8) resulted in high HR activity, indicating a strong synergistic effect. Lastly, we observed a tendency that chimeras combining W657 with two or even all three C-terminal polymorphisms (Pm3_Ch12, Pm3_Ch13 (=Pm3d), Pm3_Ch16) mount the strongest immune responses against BgtE-20069b_T86I (Fig. 6b-d, Sup Fig 14).

Next, we tested the same set of NLR chimeras against the more immune evasive variant BgtE-20069b_S74T, which is recognized by Pm3d but escapes recognition by Pm3e entirely. Again, we observed that the presence of W657 is crucial to recognize this RALPH type effector, as none of the variants lacking W657 would mount an HR response to BgtE-20069b_S74T (Fig. 6b-d, Sup Fig 14). However, in contrast to recognition of BgtE-20069b_T86I, W657 alone (Pm3_Ch2) was not sufficient for HR induction. Interestingly, the strong synergistic effect of W657 and R1356 for recognition of BgtE-20069b_T86I was also apparent for the BgtE-20069b_S74T variant, as all chimeras with this combination, namely Pm3_Ch8, Pm3_Ch13 (=Pm3d) and Pm3_Ch16 mount an efficient HR response (Fig. 6b-d, Sup Fig 14). Apart from the three chimeras with W657/R1356, only one other chimera (Pm3_Ch12: W657, R1153, V1332) was able to mount a weak HR response against BgtE-20069b_S74T. It is worth noting that Pm3d (Pm3_Ch13) exhibited the highest HR activity towards BgtE_20069b_S74T, indicating that the unique combination of polymorphisms in this NLR allele is particularly suited for recognition of BgtE-20069b variants (Fig. 6b-d, Sup Fig 14).

The unique properties of Pm3d were also apparent upon testing the set of 16 NLRs against BgtE-20069a (AvrPm3d_2). This effector escapes recognition by Pm3e entirely and results in a considerably weaker HR response upon co-expression with Pm3d compared to BgtE-20069b variants (Fig. 6ab). It therefore represents the most immune evasive variant of the RALPH-type effectors (BgtE-20069b/BgtE-20069a) tested in this context. Interestingly, among the 16 NLR chimeras only Pm3d (Pm3_Ch13) was able to mount an immune response against BgtE-20069a, again suggesting that its unique combination of polymorphisms is most suited for the recognition of the most immune evasive variants of BgtE-20069b/BgtE-20069a orthologs.

Lastly, we tested the ability of the 16 NLR chimeras to recognize the structurally divergent BgtE-5754-A1 effector (AvrPm3e_1) which is recognized by Pm3e but not by Pm3d or Pm3CS. Here, we found that the ability to recognize BgtE-5754 is largely dependent on the V1332 polymorphism found in LRR 27 of Pm3e (Pm3_Ch7). None of the chimeras carrying an E at position 1332, such as Pm3CS (Pm3_Ch1) or Pm3d (Pm3_Ch13), exhibited any HR activity towards BgtE-5754 (Fig. 6b-d, Sup Fig 14). In contrast, the introduction of V1332 alone into the Pm3CS background (Pm3_Ch4) was enough to mount an immune response, albeit at a somewhat lower level than the natural Pm3e NLR (W657 + V1332). Indeed, V1332 appeared to exhibit mild synergistic effects when combined with W657 (Pm3_Ch7 (=Pm3e), Pm3_Ch12, Pm3_Ch14, Pm3_Ch16) and was also compatible with R1153 found in Pm3d. However, we found that V1332 is not readily compatible with the other Pm3d-specific polymorphism R1356 as such combination would result in the absence of activity against BgtE-5754 (Pm3_Ch11) unless compensated by the additional presence of W657 acting synergistically with V1332 (Pm3_Ch14, Pm3_Ch16).

**Supplementary Note S7**

Combining *Pm3* alleles through conventional breeding is hampered by allelism. Furthermore, some *Pm3* alleles were found to exhibit interallelic suppression when combined[7,11,12], although interallelic suppression between *Pm3e* and *Pm3d* has not been previously assessed. We therefore crossed the highly resistant Pm3d#1 and Pm3e#2 transgenic lines and tested the resulting F1 individuals for interallelic suppression. To do so we used the isolates CHN_17-40, avirulent on *Pm3d#1* but virulent on *Pm3e#2* lines, and the isolate CHE_94202, which exhibits virulence on *Pm3d#1* but not *Pm3e#2* wheat (Fig. 7a). We found that the *Pm3d* resistance in Pm3d#1 x Pm3e#2 F1 individuals is heavily compromised, as CHN_17-40 exhibited a nearly fully virulent phenotype on these plants (Fig. 7a). Additionally, the *Pm3e* resistance was quantitatively affected in presence of *Pm3d*, as exemplified by significantly higher mildew leaf coverage of CHE_94202 on the F1 individuals as compared to the Pm3e#2 parental line (Fig. 7b). It is important to note, that in addition to interallelic suppression between *Pm3d* and *Pm3e*, also dosage effects could contribute to the observed reduction in resistance as Pm3d#1 x Pm3e#2 F1 individuals are heterozygous for both alleles, as compared to their homozygous parental lines. To further investigate the phenomenon irrespective of possible dosage effects we tested for interallelic suppression between *Pm3d* and *Pm3e* in the *N. benthamiana* system by using effector variants BgtE-20069b_S74T and BgtE-20069a, exclusively recognized by *Pm3d*, and BgtE-5754 exclusively recognized by *Pm3e*. Indeed, Pm3d-mediated HR towards BgtE-20069b_S74T or BgtE-20069a was significantly reduced in presence of Pm3e as compared to a GUS negative control (Fig. 7c). Similarly, Pm3e-mediated HR against BgtE-5754-A1 was compromised by the presence of Pm3d (Fig. 7c). Taken together, our results confirm the occurrence of interallelic suppression between *Pm3d* and *Pm3e* and suggests that stacking these two alleles in the same wheat line would not result in an additive effect.

### Supplementary Figures

A

| | Signalp 4.1 | | Signalp 5.0 | | | Signalp 6.0 | | | DeepTHMM | |
|---|---|---|---|---|---|---|---|---|---|---|
| Gene | Predicted cleavage side | D-score (Cutoff > 0.450) | Probablity signal peptide | Cleavage side | Probability cleavage side | Probablity signal peptide | Cleavage side | Probability cleavage side | Most likely predicted topology | Cleavage side |
| BgtE-5754 | 16/17 | 0,476 | 0,611 | 15/16 | 0,170 | 0,426 | 16/17 | 0,574 | Globular & SP | 13/14 |
| BgtE-20069b | 21/22 | 0,923 | 0,990 | 21/22 | 0,939 | 1,000 | 21/22 | 0,979 | Globular & SP | 21/22 |

B

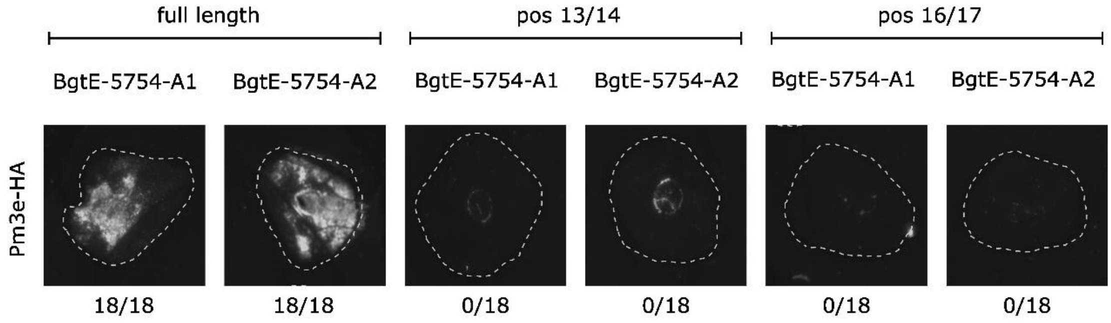


**Supplementary Figure 1. The full-length BgtE-5754 effector is recognized by Pm3e. A)** Summary of signal peptide prediction of BgtE-5754 and BgtE-20069b with various prediction software. **B)** Co-infiltration of Pm3e-HA with various BgtE-5754 truncation variants in *N. benthamiana*. "Full length" refers to the use of the complete coding sequence (CDS), whereas "pos 13/14" and "pos 16/17" indicate truncated versions of the CDS corresponding to the predicted cleavage sites of a potential signal peptide at those respective positions in the mature protein. Co-expression was performed using a 4 (effector) : 1 (NLR) : 1 (p19 silencing inhibitor) ratio and imaged with a Fusion FX imager. The experiment was performed three times with a total of n=18 replicates. A representative picture of the HR response is shown. The number of replicates with a detectable HR response are indicated below the images.

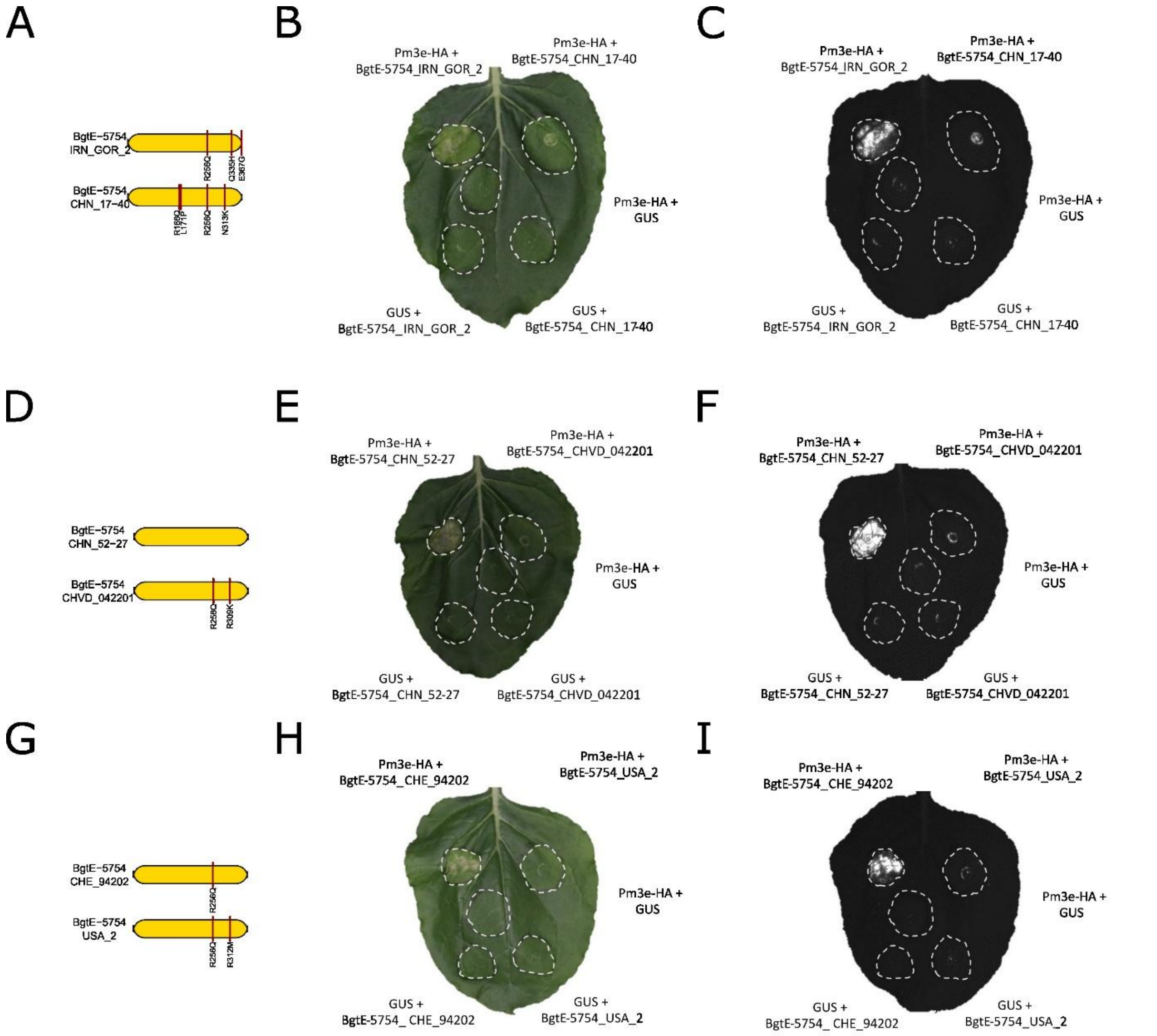


**Supplementary Figure 2. Analysis of BgtE-5754 variants in the parental isolates of three genetic crosses that identify the *AvrPm3e_1* locus. A)** Protein model of the AvrPm3e_1 candidate effector BgtE-5754 in the *Pm3e*-avirulent isolate IRN_GOR_2 and the virulent isolate CHN_17-40, with non-synonymous polymorphisms indicated by red lines in the protein, compared to the reference sequence BgtE-5754-A1. **B & C)** *Agrobacterium*-mediated co-expression of BgtE-5754_IRN_GOR_2 (BgtE-5754-A2) or BgtE-5754_CHN_17-40 (BgtE-5754-V1) with Pm3e-HA or a GUS negative control in *N. benthamiana*. **D)** Protein model of the AvrPm3e_1 candidate effector BgtE-5754 in the *Pm3e*-avirulent isolate CHE_94202 and the *Pm3e*-avirulent isolate USA_2, with non-synonymous polymorphisms in the protein indicated by red lines compared to the reference sequence BgtE-5754-A1. **E & F)** *Agrobacterium*-mediated co-expression of BgtE-5754_CHE_94202 (BgtE-5754-A3) or BgtE-5754_USA_2 (BgtE-5754-V3) with Pm3e-HA or a GUS negative control in *N. benthamiana*. **G)** Protein model of the AvrPm3e_1 candidate effector BgtE-5754 in the *Pm3e*-avirulent isolate CHN_52-27 and the *Pm3e*-virulent isolate CHVD_042201, with non-synonymous polymorphisms in the protein indicated by red lines compared to the reference sequence BgtE-5754-A1.**H & I)** *Agrobacterium*-mediated co-expression of BgtE-5754_CHN_52-27 (BgtE-5754-A1) or BgtE-5754_CHVD_042201 (BgtE-5754-V3) with Pm3e-HA or a GUS negative control in *N. benthamiana*. Co-expressions were performed using a 4 (effector) : 1 (NLR) : 1 (p19 silencing inhibitor) ratio and imaged either with a standard camera (B, E, H) or a Fusion FX imager (C, F, I). All experiments were repeated three times with n = 6 leaves per experiment (total n = 18).

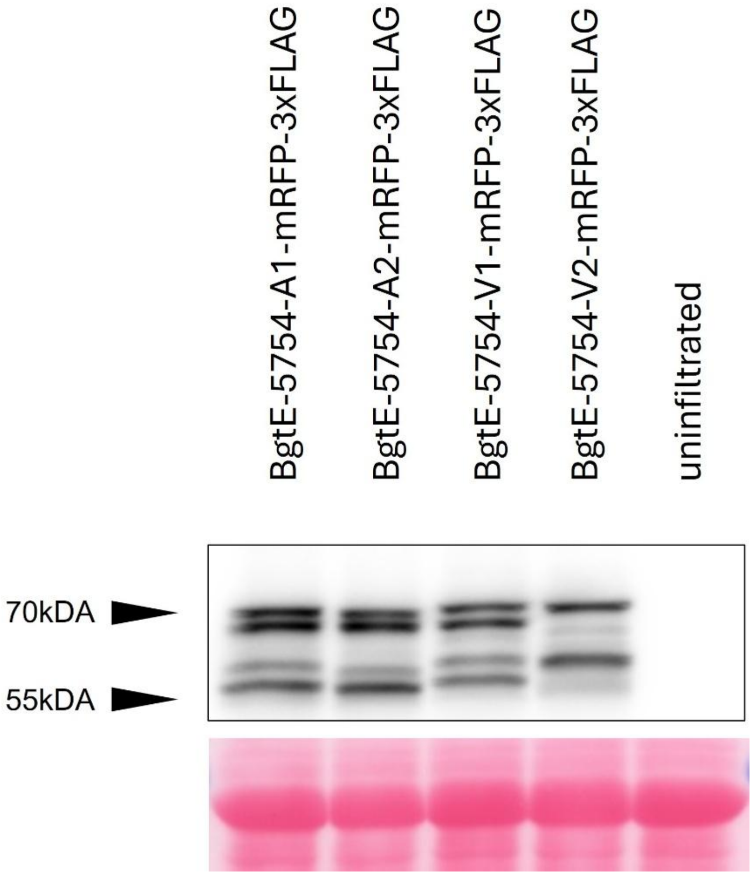


**Supplementary Figure 3. Western blot analysis of selected BgtE-5754 variants.** BgtE-5754-mRFP-3xFLAG variants A1, A2, V1 and V2 were expressed in *N. benthamiana* and detected via anti-FLAG western blotting. Proteins obtained from uninfiltrated tissue were included as negative control. Ponceau S staining of the blot was used to ensure equal loading and is shown in the lower panel. The experiment was performed three times with similar results.

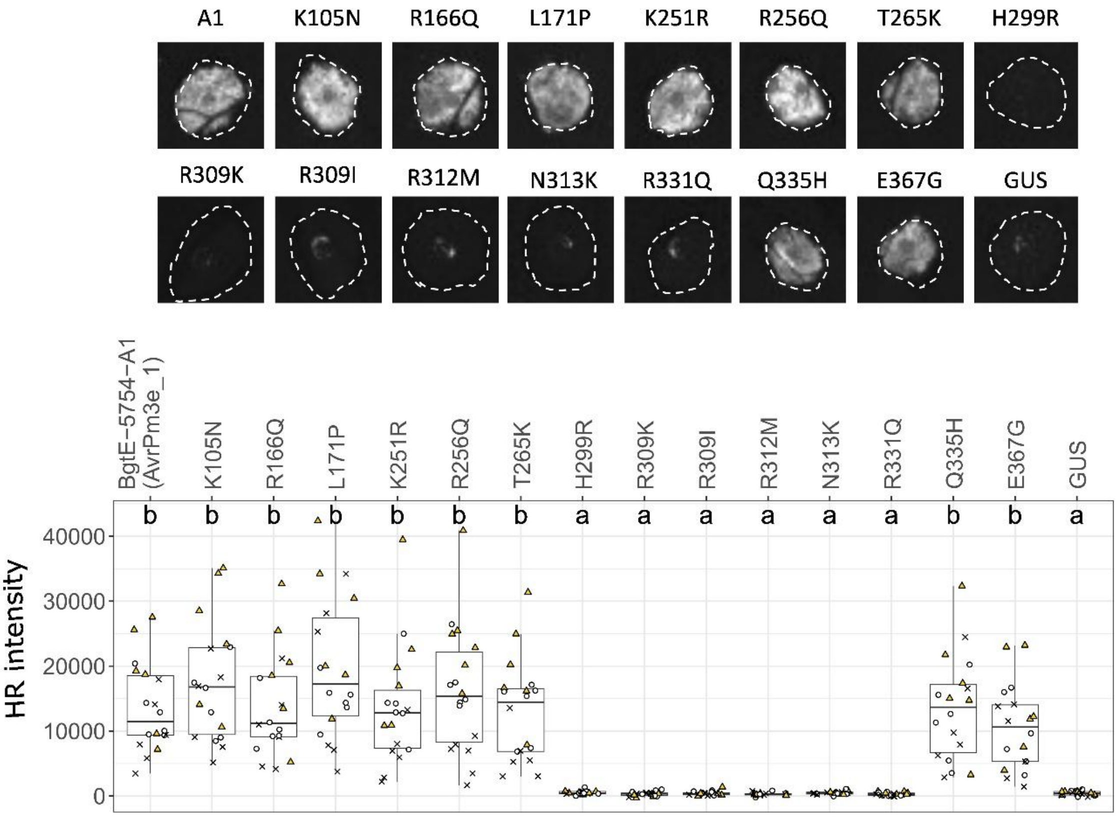


**Supplementary Figure 4. Effect of individual amino acid substitutions in BgtE-5754-A1 on the recognition by Pm3e**. Each amino acid substitution was introduced separately into the BgtE-5754-A1 background and tested for recognition by Pm3e-HA via *Agrobacterium*-mediated co-expression in *N. benthamiana*. Co-expression was performed using a 4 (effector) : 1 (NLR) : 1 (p19 silencing inhibitor) ratio and imaged with a Fusion FX imager. Top panel shows representative HR pictures for each combination. Lower panel displays quantified HR intensities from 18 individual infiltration spots per combination, collected across three independent experiments (indicated by different symbols). Letters above boxplots denote statistically significant differences at $p < 0.05$ based on Dunn's test.

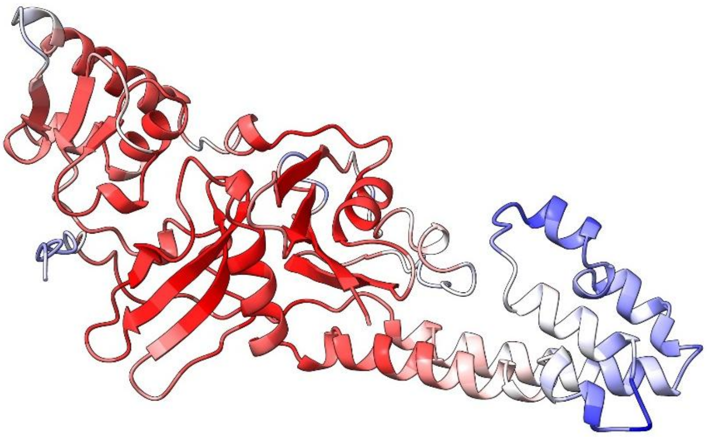

**Supplementary Figure 5. Predicted 3D structure of BgtE-5754 according to Alphafold3 modelling.** Coloring represents predicted local distance difference test (pLDDT) local confidence according to Alphafold (100 (dark red) – 0 (dark blue)).

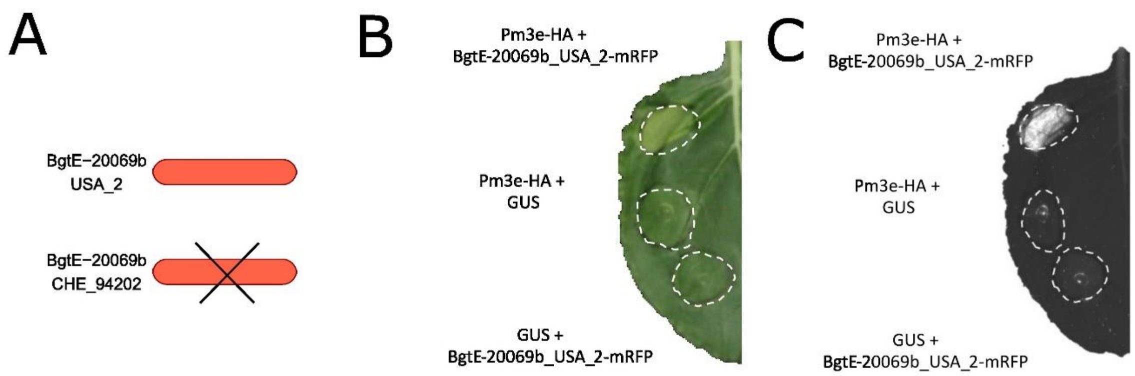


**Supplementary Figure 6. Characterization of BgtE-20069b variants found in the parental isolates of cross USA_2 x CHE_94202. (A)** Gene models of *BgtE-20069b* in the *Pm3e*-avirulent isolate USA_2 and the *Pm3e*-avirulent isolate CHE_94202. The deletion of *BgtE-20069b* in CHE_94202 is indicated by crossed black lines. **(B & C)** *Agrobacterium*-mediated co-expression of BgtE-20069b-mRFP (AvrPm3e_2) with Pm3e-HA or a GUS negative control in *N. benthamiana*. Co-expression was performed using a 4 (effector) : 1 (NLR) : 1 (p19 silencing inhibitor) ratio and imaged with a camera (B) or a Fusion FX imager (C). The experiment was performed three times with similar results with n=6 leaves per experiment (total n=18).

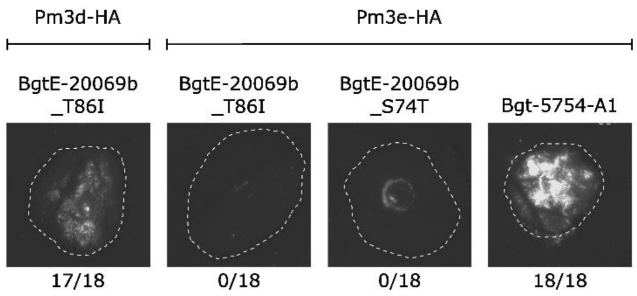


**Supplementary Figure 7. BgtE-20069b_T86I without mRFP stabilization does not induce HR upon co-expression with Pm3e in *N. benthamiana*.** Untagged BgtE-20069b_T86I was co-expressed with either Pm3d-HA or Pm3e-HA. BgtE-5754-A1 co-expression with Pm3e-HA was included as a positive control. Co-expression was performed using a 4 (effector) : 1 (NLR) : 1 (p19 silencing inhibitor) ratio and imaged with a Fusion FX imager The experiment was performed three times with similar results with n=6 leaves per experiment (total n=18). A representative picture of the HR response is shown. The number of replicates with a detectable HR response are indicated below the image.

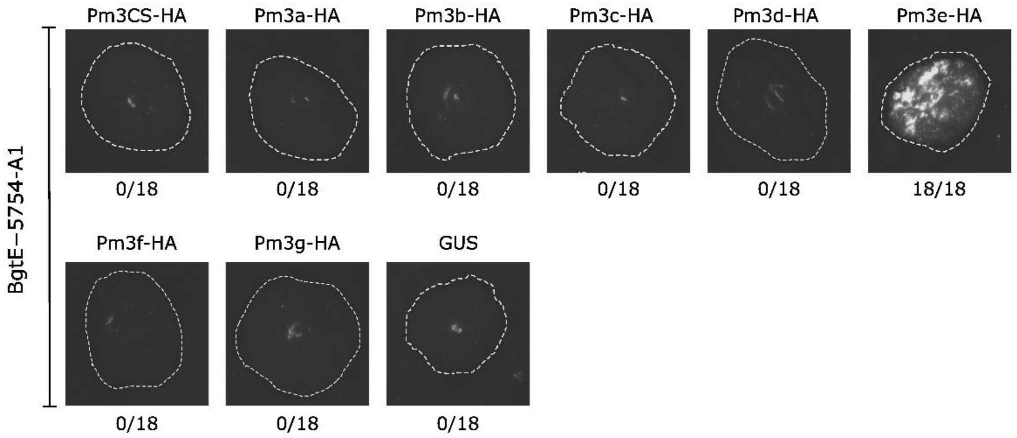


**Supplementary Figure 8. BgtE-5754 is not recognized by other members of the *Pm3* allelic series (*Pm3a-g*).** BgtE-5754-A1 was co-expressed with C-terminally HA-tagged Pm3CS, Pm3a, Pm3b, Pm3c, Pm3d, Pm3e, Pm3f, Pm3g or a GUS negative control in *N. benthamiana*. Co-expression was performed using a 4 (effector) : 1 (NLR) : 1 (p19 silencing inhibitor) ratio and imaged with a Fusion FX imager. Each combination was tested 18 times (n=18) in three independent experiments. A representative picture of the HR response is shown. The number of replicates with a detectable HR response are indicated below the image.

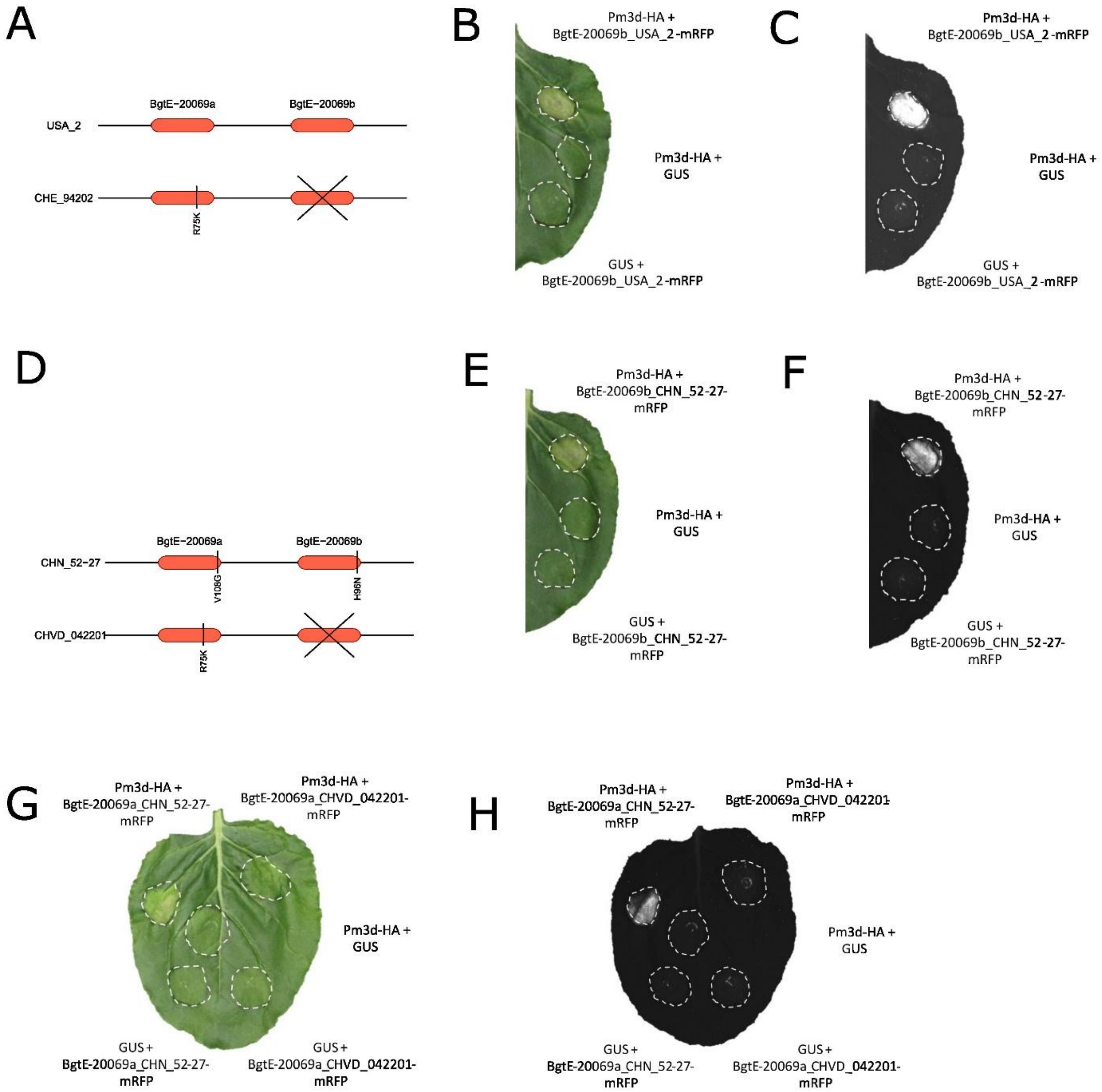


**Supplementary Figure 9. Characterization of BgtE-20069b (AvrPm3d_1) and BgtE-20069a (AvrPm3d_2) variants in the parental isolates of crosses USA_2 x CHE_94202 and CHN_52-27 x CHVD_042201. (A)** Gene models of *BgtE-20069a* and *BgtE-20069b* in the *Pm3d*-avirulent isolate USA_2 and the *Pm3d*-virulent isolate CHE_94202. The deletion of *BgtE-20069b* in CHE_94202 is indicated by crossed black lines. **(B & C)** *Agrobacterium*-mediated co-expression of BgtE-20069b-mRFP (AvrPm3d_1) with Pm3d-HA or a GUS negative control in *N. benthamiana*. **(D)** Gene models of *BgtE-20069a* and *BgtE-20069b* in the *Pm3d*-avirulent isolate CHN_52-27and the *Pm3d*-virulent isolate CHVD_042201. The deletion of *BgtE-20069b* in CHVD_042201 is indicated by crossed black lines. **(E & F)** *Agrobacterium*-mediated co-expression of BgtE-20069b_H96N-mRFP (AvrPm3d_1) with Pm3d-HA or a GUS negative control in *N. benthamiana*. **(G & H)** *Agrobacterium*-mediated co-expression of BgtE-20069a_V108G-mRFP (AvrPm3d_2) and BgtE-20069a_R75K-mRFP with Pm3d-HA or a GUS negative control in *N. benthamiana*.

All co-expression experiments were performed using a 4 (effector) : 1 (NLR) : 1 (p19 silencing inhibitor) ratio and imaged with a camera (**B,E,G**) or a Fusion FX imager (**C,F,H**). The experiment was performed three times with similar results with n=6 leaves per experiment (total n=18).

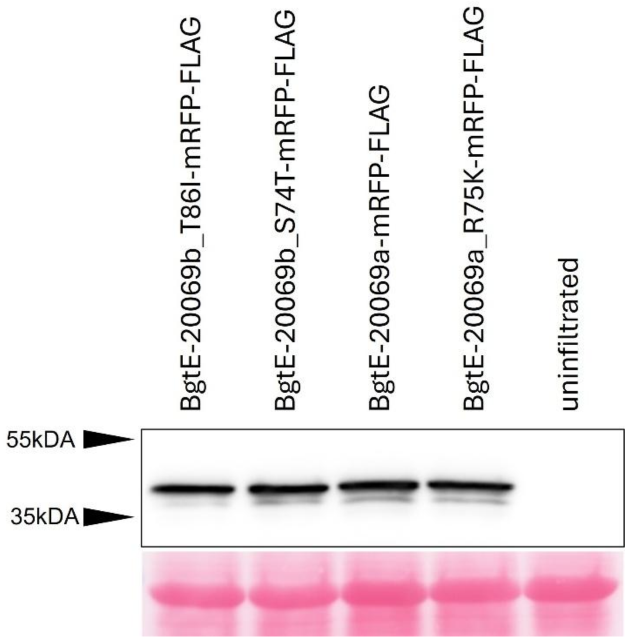


**Supplementary Figure 10. Western blot analysis of selected BgtE-20069b/BgtE-20069a variants.** BgtE-20069b-mRFP-FLAG and BgtE-20069a-mRFP-FLAG variants were expressed in *N. benthamiana* and detected via anti-FLAG western blotting. Proteins obtained from uninfiltrated tissue were included as negative control. Ponceau S staining of the blot was used to ensure equal loading and is shown in the lower panel. The experiment was performed three times with similar results.

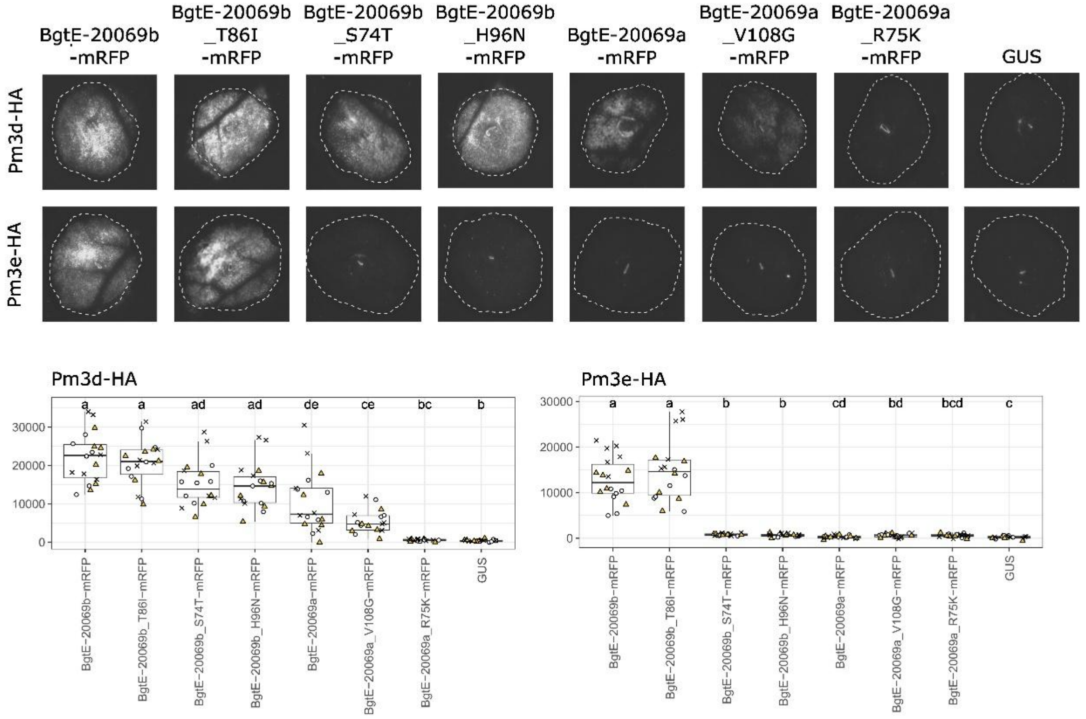


**Supplementary Figure 11. Recognition of various BgtE-20069b and BgtE-20069a variants by Pm3d and Pm3e.** C-terminally tagged mRFP variants of BgtE-20069b and BgtE-20069a were tested for recognition by Pm3d-HA and Pm3e-HA via *Agrobacterium*-mediated co-expression in *N. benthamiana*. Co-expression was performed using a 4 (effector) : 1 (NLR) : 1 (p19 silencing inhibitor) ratio and imaged with a Fusion FX imager. The top panel shows representative images for each combination. Bottom panel displays quantified HR intensities from 18 individual infiltration spots per combination, collected across three independent experiments (indicated by different symbols). Letters above boxplots denote statistically significant differences at $p < 0.05$ based on Dunn's test.

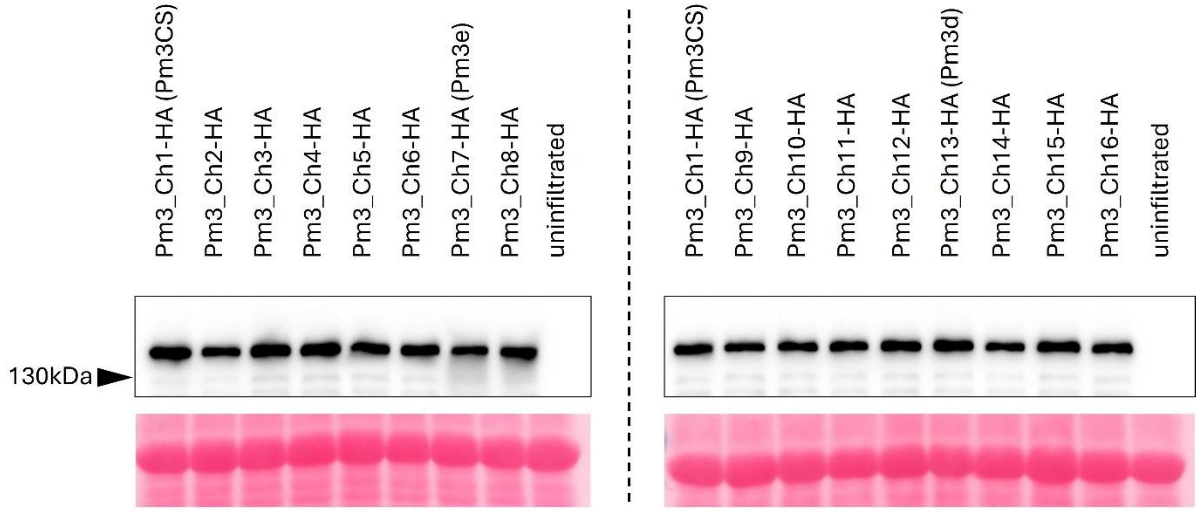


**Supplementary Figure 12. Western blot analysis of the 16 chimeric C-terminally HA-tagged Pm3 proteins.** Proteins were analyzed on two separate western blots, as indicated by the dashed line. In each western blot Pm3_Ch1-HA (Pm3CS) is shown for comparison and protein extracts from non-infiltrated tissue serve as negative controls. Ponceau S staining of the blot was used to ensure equal loading and is shown in the lower panel. The experiment was performed three times with similar results.

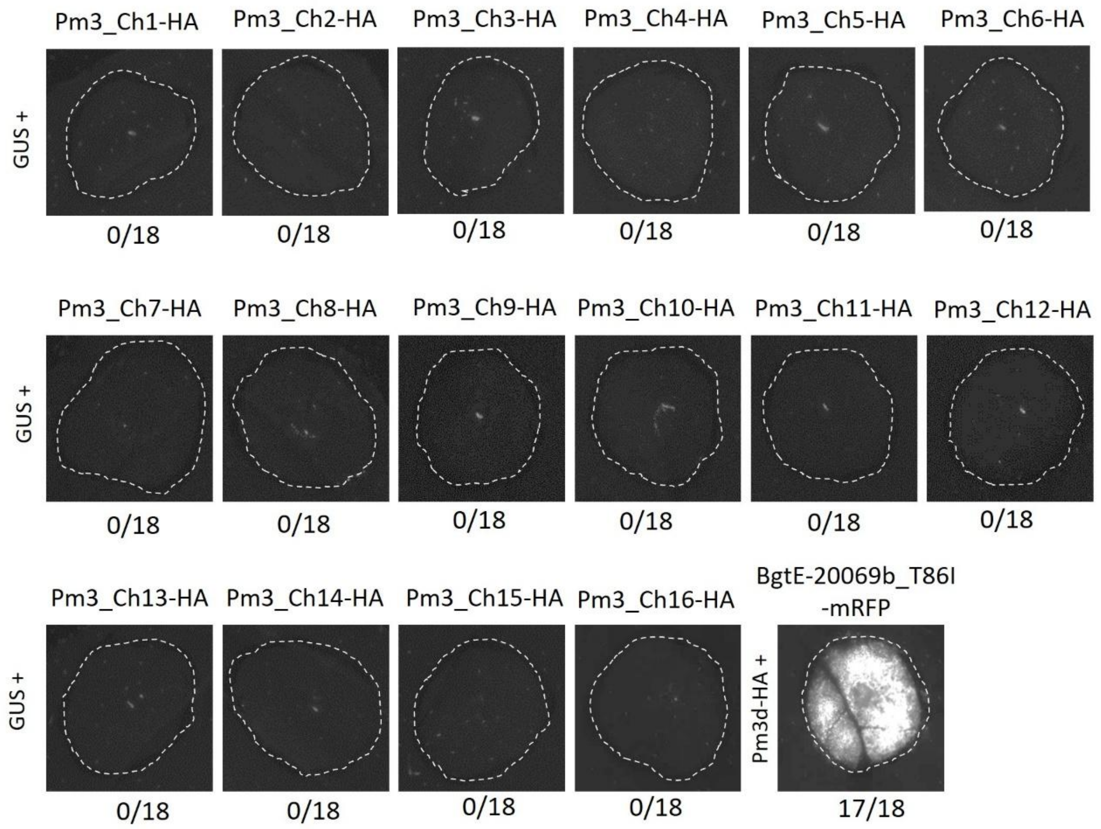


**Supplementary Figure 13. Chimeric Pm3 constructs (Pm3_Ch1-16) do not show autoactivity upon expression in *N. benthamiana*.** C-terminally HA-tagged chimeric Pm3 constructs were co-expressed with GUS via *Agrobacterium*-mediated transformation. Co-expression was performed using a 4 (GUS) : 1 (NLR) : 1 (p19 silencing inhibitor) ratio and imaged with a Fusion FX imager. Co-expression of BgtE-20069b_T86I-mRFP with Pm3d-HA in a 4 (AVR) : 1 (NLR) : 1 (p19 silencing inhibitor) ratio was used as a control. Each combination was tested 18 times (n=18) in three independent experiments. A representative picture of the HR response is shown. The number of replicates with a detectable HR response are indicated below the image.

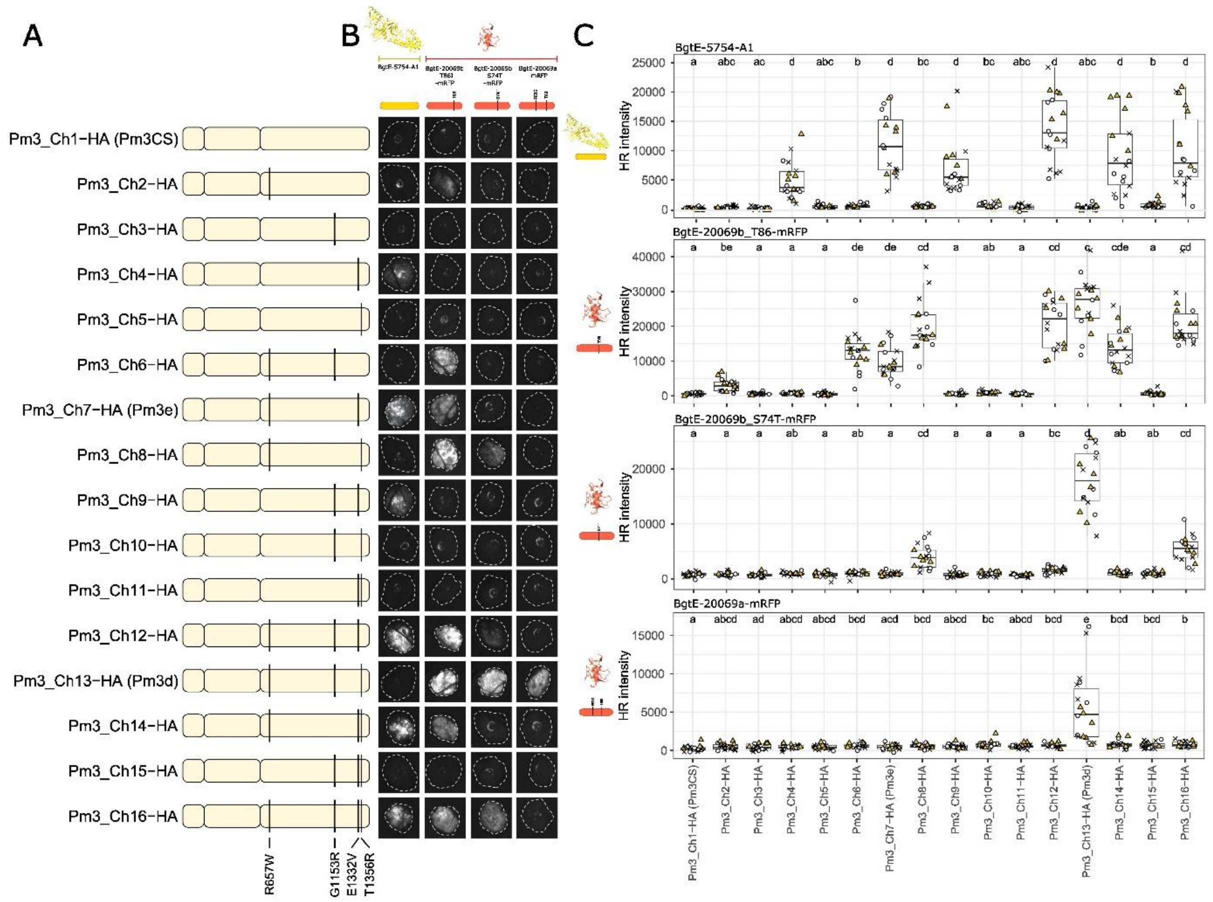


**Supplementary Figure 14. Pm3d and Pm3e NLR specificity is controlled by individual amino acid polymorphisms in the LRR domain (A)** Schematic representation of 16 chimeric NLR variants used to study Pm3d and Pm3e specificity. CC-, NB-ARC- and LRR domains are represented by light yellow boxes (from left to right). Pm3CS is shown as a reference. The positions of polymorphic residues are indicated by black lines. **(B)** *Agrobacterium*-mediated co-expression of BgtE-5754, BgtE-20069b_T86I-mRFP, BgtE-20069_S74T-mRFP or BgtE-20069a-mRFP with each of the C-terminally HA-tagged 16 chimeric NLR variants in *N. benthamiana*. Co-expression was performed using a 4 (effector) : 1 (NLR) : 1 (p19) ratio and imaged with a Fusion FX imaging system. The experiment was performed three times with n=6 leaves per experiment (total n=18). **(C)** Quantification of the HR response depicted in (B). Datapoints from the three independent experiments are color coded and represented by different symbols (cross, triangle, circle). Letters above the individual boxplots represent statistical differences according to Dunn's test ($p<0.05$).

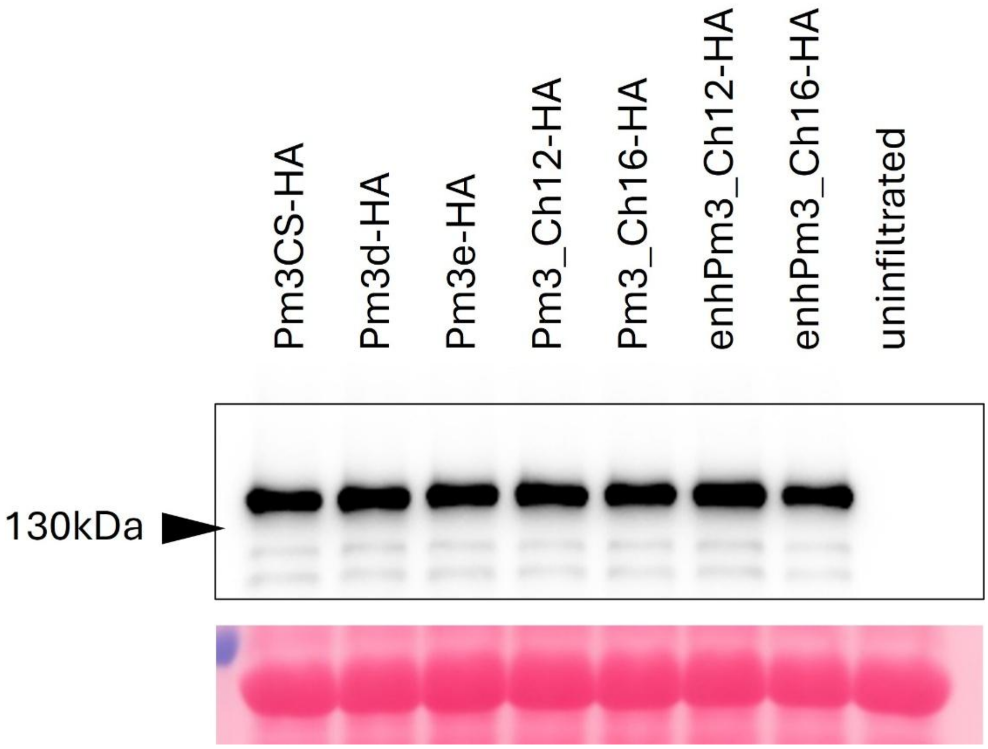


**Supplementary Figure 15. Western blot analysis of C-terminally HA-tagged chimeric Pm3 proteins.** Protein extracts from non-infiltrated *N. benthamiana* tissue served as a negative control. Ponceau S staining of the blot was used to ensure equal loading and is shown in the lower panel. The experiment was performed three times with similar results.

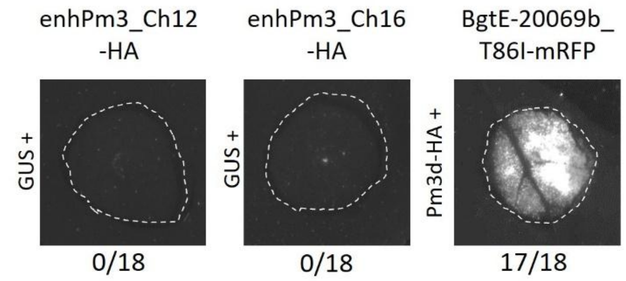


**Supplementary Figure 16. Enhanced chimeric Pm3 constructs enhPm3_12 and enhPm3_Ch16 do not show autoactivity upon expression in *N. benthamiana*.** The C-terminally HA-tagged enhPm3_Ch12 and enhPm3_Ch16 constructs were co-expressed with GUS via *Agrobacterium*-mediated transformation in *N. benthamiana*. Co-expression was performed using a 4 (GUS) : 1 (NLR) : 1 (p19 silencing inhibitor) ratio and imaged with a Fusion FX imager. Co-expression of BgtE-20069b_T86I-mRFP with Pm3d-HA in a 4 (AVR) : 1 (NLR) : 1 (p19 silencing inhibitor) ratio is shown as a control. Each combination was tested 18 times (n=18) in three independent experiments. A representative picture of the HR response is shown. The number of replicates with a detectable HR response are indicated below the image.

**Supplementary Table 1: Summary of the identified intervals using the avirulence depletion assay in different *Bgt* crosses**

| *Bgt* cross | Selection line | Reference assembly | Chromosome | Interval positions |
|---|---|---|---|---|
| CHN_52-27 X CHN_17-40 | Pm3e#2 | Bgt_CHN_52_27_genome_v1 | Chr-04 | 3157657,3192884 |
| CHN_52-27 X CHN_17-40 | Pm3e#2 | Bgt_genome_v3_16 | Chr-04 | 3135295,3180690 |
| CHN_52-27 X CHVD_042201 | Pm3e#2 | Bgt_CHN_52_27_genome_v1 | Chr-04 | 2954498,3174843 |
| CHN_52-27 X CHVD_042201 | Pm3e#2 | Bgt_genome_v3_16 | Chr-04 | 2938459,3157602 |
| IRN_GOR_2 X CHN_17-40 | Pm3e#2 | Bgt_IRN_GOR_2_genome_v1 | Chr-04 | 2974526,3206780 |
| IRN_GOR_2 X CHN_17-40 | Pm3e#2 | Bgt_genome_v3_16 | Chr-04 | 2929978,3138563 |
| IRN_GOR_2 X CHN_17-40 | Pm3e#2 | Bgt_IRN_GOR_2_genome_v1 | Chr-09 | 394040,826195 |
| IRN_GOR_2 X CHN_17-40 | Pm3e#2 | Bgt_genome_v3_16 | Chr-09 | 334180,646221 |
| USA_2 X CHE_94202 | Pm3e#2 | Bgt_USA_2_genome_v1 | Chr-04 | 2525685,3585533 |
| USA_2 X CHE_94202 | Pm3e#2 | Bgt_genome_v3_16 | Chr-04 | 2454741,3377968 |
| USA_2 X CHE_94202 | Pm3e#2 | Bgt_USA_2_genome_v1 | Chr-09 | 369147,775630 |
| USA_2 X CHE_94202 | Pm3e#2 | Bgt_genome_v3_16 | Chr-09 | 294502,686545 |
| CHN_52-27 X CHVD_042201 | Pm3d#1 | Bgt_CHN_52_27_genome_v1 | Chr-09 | 353877,765980 |
| CHN_52-27 X CHVD_042201 | Pm3d#1 | Bgt_genome_v3_16 | Chr-09 | 296970,750271 |
| USA_2 X CHE_94202 | Pm3d#1 | Bgt_USA_2_genome_v1 | Chr-09 | 311508,828481 |
| USA_2 X CHE_94202 | Pm3d#1 | Bgt_genome_v3_16 | Chr-09 | 242992,772390 |

**Supplementary Table 2: Normalized read coverage in isolate CHE_96224 and its derived UV-mutant Bgt_FAB**

| Gene | CHE_96224 | Bgt_FAB |
|---|---|---|
| *BgtE-20069a* | 0,98 | 1,16 |
| *BgtE-20069b* | 1,10 | 0,13 |
| *Bgt-712* (*GAPDH*) | 0,86 | 0,99 |
| *BgtE-5754* (*AvrPm3e_1*) | 0,94 | 0,93 |

**Supplementary Table 3: Summary of isolates that carry a deletion in the *BgtE-20069b* gene**

| Isolate | Normalized coverage *BgtE-20069a* | Normalized coverage *BgtE-20069b*[1] | Genotype *BgtE-20069a* | Genotype *BgtE-20069b* | Phenotype Pm3d#1[2] |
|---|---|---|---|---|---|
| FRA_SYROS_2000_15 | 0,819 | 0,108 | BgtE-20069a_R75K | Deletion | NA |
| FRA_SYROS_05_54 | 0,840 | 0,115 | BgtE-20069a_R75K | Deletion | NA |
| GBR_WC1110 | 0,767 | 0,116 | BgtE-20069a | Deletion | I |
| ISR_K_U | 0,893 | 0,117 | BgtE-20069a | Deletion | I |
| CHE_96229 | 0,959 | 0,120 | BgtE-20069a_R75K | Deletion | V |
| USA_KAN_43 | 1,005 | 0,121 | BgtE-20069a | Deletion | NA |
| CHE_10001 | 0,936 | 0,122 | BgtE-20069a_R75K | Deletion | NA |
| CHE_07298 | 0,992 | 0,134 | BgtE-20069a_R75K | Deletion | V |
| GBR_JIW48 | 1,835 | 0,167 | 2 x BgtE-20069a | Deletion | A |
| CHE_94202 | 0,727 | 0,180 | BgtE-20069a_R75K | Deletion | V |
| CHE_98013 | 0,830 | 0,188 | BgtE-20069a_R75K | Deletion | V |
| CHN_5_93 | 1,470 | 0,379 | 2 x BgtE-20069a | Deletion | A |

[1] Isolates with a normalized mapping coverage of <0.4 over the coding sequence of BgtE-20069b are listed

[2] Virulence phenotypes of tested isolates on the Pm3d#1 transgenic line. Phenotype categories include: A = avirulent (0% mildew leaf coverage), I = intermediate (20-50% mildew leaf coverage), V = virulent (100% leaf coverage), NA = not available

**Supplementary Table 4: List of primers used in this study**

| Name | Sequence (5'-3') | Purpose |
|---|---|---|
| LK315 | CGTCATTTGAAGAGACTTGTG | haplovariant mining of *BgtE-20069a* and *BgtE-20069b* |
| LK316 | CTAATCTCGACAACTCTGTTATCG | haplovariant mining of *BgtE-20069a* and *BgtE-20069b* |
| LK994 | GATCTGAAAATTCGCGACAAAAAGC | haplovariant mining of *BgtE-5754* |
| LK996 | CATACAGCTATGCCCAACACAC | haplovariant mining of *BgtE-5754* |
| LK982 | CACTCATGGATGTTGGAACTTGAAG | SDM primer to introduce W657 polymorphism in Pm3CS/Pm3d/Pm3e LRR |
| LK983 | CACTCATGGATGTCGGAACTTGAAG | SDM primer to introduce R657 polymorphism in Pm3CS/Pm3d/Pm3e LRR |
| LK984 | TAGAGGTGGCAGAGGGAAG | reverse primer to combine with LK982 or LK983 |
| LK985 | AAAATGACTATTCGTGGGTGCATTAAG | SDM primer to introduce R1153 polymorphism in Pm3CS/Pm3d/Pm3e LRR |
| LK986 | AAAATGACTATTGGTGGGTGCATTAAG | SDM primer to introduce G1153 polymorphism in Pm3CS/Pm3d/Pm3e LRR |
| LK987 | CTTGAGAGATGCCGGGAC | reverse primer to combine with LK985 or LK986 |
| LK988 | CTTTGGCTTGAAAGATGCAGTACC | SDM primer to introduce E1332 polymorphism in Pm3CS/Pm3d/Pm3e LRR |
| LK989 | CTTTGGCTTGTAAGATGCAGTACC | SDM primer to introduce V1332 polymorphism in Pm3CS/Pm3d/Pm3e LRR |
| LK990 | GGATTCCAGCGATGGGG | reverse primer to combine with LK988 or LK989 |
| LK991 | CTTGAAATTAGAGGCTGCCCTG | SDM primer to introduce R1356 polymorphism in Pm3CS/Pm3d/Pm3e LRR |
| LK992 | CTTGAAATTACAGGCTGCCCTG | SDM primer to introduce T1356 polymorphism in Pm3CS/Pm3d/Pm3e LRR |
| LK993 | AGACCAGAGAGACCTGTATACTTG | reverse primer to combine with LK991 or LK992 |
| LK1011 | AACACAAGGAAGATAGTCCCGAAACA | SDM primer to introduce L456P/Y458H polymorphisms into Pm3 NB-ARC |
| LK1012 | CAGGGATAAAGCCGTTTGCAATC | SDM primer to introduce L456P/Y458H polymorphisms into Pm3 NB-ARC |
